# Local structural dynamics of alpha-synuclein correlate with aggregation in different physiological conditions

**DOI:** 10.1101/2022.02.11.480045

**Authors:** Neeleema Seetaloo, Maria Zacharopoulou, Amberley D. Stephens, Gabriele S. Kaminski Schierle, Jonathan J. Phillips

## Abstract

In Parkinson’s disease and other synucleinopathies, the intrinsically disordered, presynaptic protein alpha-synuclein misfolds and aggregates. We hypothesise that the exposure of alpha-synuclein to different cellular environments, with different chemical compositions, pH and binding partners, alters its biological and pathological function by inducing changes in molecular conformation. Our custom instrumentation and software enable measurement of the amide hydrogen exchange rates of wild-type alpha-synuclein at amino acid resolution under physiological conditions, mimicking those in the extracellular, intracellular, and lysosomal compartments of cells. We characterised the aggregation kinetics and morphology of the resulting fibrils and correlate these with structural changes in the monomer. Our findings reveal that the C-terminal residues of alpha-synuclein are driving its nucleation and thus its aggregation. Furthermore, the entire NAC region and specific other residues strongly promoted elongation of fibrils. This provides new detail on our current understanding of the relationship between the local chemical environment and monomeric conformations of alpha-synuclein.

## INTRODUCTION

Parkinson’s disease (PD) is a neurodegenerative condition affecting over 6.2 million people worldwide and this number is predicted to reach 13 million by 2040^1^. One of the hallmarks of PD is the appearance of cytoplasmic inclusions in neurons, known as Lewy bodies and Lewy neurites, which are mostly constituted of β-sheet-rich aggregates of the protein alpha-synuclein (aSyn)^2^. Being intrinsically disordered, monomeric aSyn exists as an ensemble of interconverting protein structures^3^. In PD and other synucleinopathies, soluble disordered monomeric aSyn can misfold and aggregate, first forming oligomeric species before culminating to insoluble, highly structured amyloid fibrils^4^. aSyn is a 14.46 kDa protein consisting of 140 amino acid residues divided into three domains: a highly positively charged amphipathic N-terminus (1-60), a central hydrophobic core (61-95) known as the non-amyloid beta component (NAC), and an acidic C-terminal tail (96-140) (Supplementary Figure 1). Unlike well-folded proteins, being natively unfolded, aSyn adopts a broad but shallow conformational space, meaning it can interchange with other conformers with minimal activation energy^5^. Using a variety of techniques including nuclear magnetic resonance and mass spectrometry, it has been found that the conformations adopted by monomeric aSyn are stabilised by long-range intramolecular electrostatic and hydrophobic interactions between its charged N- and C-termini, and between the C-terminus and the NAC region^6–9^. These intramolecular interactions account for its smaller radius of gyration, compared to the prediction of a 140-residue protein random coil, suggesting a partially-folded structure^10^. Disruptions in these long-range interactions, such as mutations, changes in the local environments and post-translational modifications (PTMs), can skew the conformational ensemble and disturb the stability of the protein, inducing misfolding and aggregation^11^. Therefore, it remains crucial to establish the correlation between monomeric conformation and aggregation propensity/kinetics of aSyn.

Whilst it has been found to be widely distributed in the body^12^, aSyn is particularly enriched at the presynapse (∼20-40 µM)^13^ and has been proposed to participate in the homeostasis and recycling of synaptic vesicles^14^. aSyn encounters a number of different cellular and extracellular environments through various routes (summarised in Table 1^15^): (i) exposure to the extracellular space via exocytosis, apoptosis, exosome release, and release of cellular contents^16^; (ii) endocytosis into the endosomal/lysosomal pathway^17^; (iii) Metabolic imbalances leading to calcium and mitochondrial dysfunction^18,19^. aSyn in these different chemical environments will have a uniquely biased conformational ensemble^20,21^. This leads to the crucial question of whether these different conformational ensembles in the monomer correlate with the propensity and kinetics of aggregation. Furthermore, these differences in structural dynamics of the monomer may result in different fibril morphologies, which could be indicative of alternative aggregation mechanisms.

**Table 1:** Composition of extracellular, intracellular, and lysosomal compartments used in this study (adapted from Stephens et al^15^). The maximum potential concentration was used for all ions.

| Ion | Extracellular<br>concentration (mM) | Intracellular<br>concentration (mM) | Lysosomal<br>concentration (mM) |
| --- | --- | --- | --- |
| Na<sup>+</sup> | 143 | 15 | 20 |
| K<sup>+</sup> | 4 | 140 | 60 |
| Ca<sup>2+</sup> | 2.5 | 100 nM | 0<sup>a</sup> |
| Mg<sup>2+</sup> | 0.7 | 10 | 0 |
| pH | 7.4 | 7.2 | 4.9 |
| Buffer<br>system | 20 mM Tris | 20 mM Tris | 20 mM citrate |
| References<sup>b</sup> | 110746 BNID<sup>36</sup> | <sup>36</sup>103966, 110746 BNID | <sup>37,38</sup>114028 BNID |

<sup>a</sup>Conflicting reports as to whether lysosomes are Ca<sup>2+</sup> stores (0.5-0.6 mM)<sup>39</sup>.
<sup>b</sup>BNID numbers correspond to The Database of Useful Biological Numbers; http://bionumbers.hms.harvard.edu/search.aspx.

Furthermore, a variety of oligomer species^22^ and fibril polymorphs^23^ have been discovered, with different biophysical properties and levels of toxicity^24–26^, possibly dependent on the local environments in which they arise, familial mutations^27–32^, or PTMs^33^. There is growing evidence supporting the idea that the monomeric aSyn conformation and the surrounding environment affects fibril formation^34,35^. Here, we aim to understand whether and how conformational changes in the monomer may affect the aggregation kinetics and fibril morphologies across different solution conditions mimicking cellular and extracellular compartments in vitro.

Here, we obtained data at high structural and temporal resolution for the aSyn monomer under physiologically relevant conditions. This was achieved by hydrogen-deuterium exchange mass spectrometry (HDX-MS) on the millisecond timescale coupled with a gas-phase ‘soft fragmentation’ technique known as electron-transfer dissociation (ETD). We correlated these data to Thioflavin-T (ThT-) based aggregation kinetics and fibril morphology, assessed by atomic force microscopy (AFM). Our results show that the solution conditions assessed in this study all lead to distinct aggregation kinetics, fibril morphologies and monomeric conformations. More importantly, our correlative analyses reveal specific local conformational changes in the aSyn monomer that influence the separate stages of aggregation, namely the nucleation and elongation steps.

## RESULTS

### Aggregation propensity increases from Tris-only < Extracellular < Intracellular < Lysosomal conditions *in vitro*

We first investigated whether the aggregation propensity of aSyn differed across four conditions with varying pH and ionic compositions, mimicking the extracellular, intracellular and lysosomal environments, alongside our baseline Tris-only condition (20 mM Tris, pH 7.4). To do so, we used a ThT-based fluorescence assay. The ThT molecule emits fluorescence when bound to rich fibrillar β-sheet structures, informing us on the process of aggregation^40^ (Figure 1A, 1C-1F). In the ThT-based assay, the time before the onset of fluorescence, lag time (t_*lag*_), is indicative of the nucleation phase of fibril formation and the slope of the exponential growth (k_*agg*_) describes the elongation phase (Figure 1B). Upon the addition of physiologically relevant salts, the aSyn monomer nucleation lag time is reduced by 45% from 113 h to 62 h and the elongation rate k_agg_ is increased by 42% (0.024 h^-1^ to 0.034 h^-1^), as can be seen from Figure 1C and 1E, respectively.

**Figure 1:**
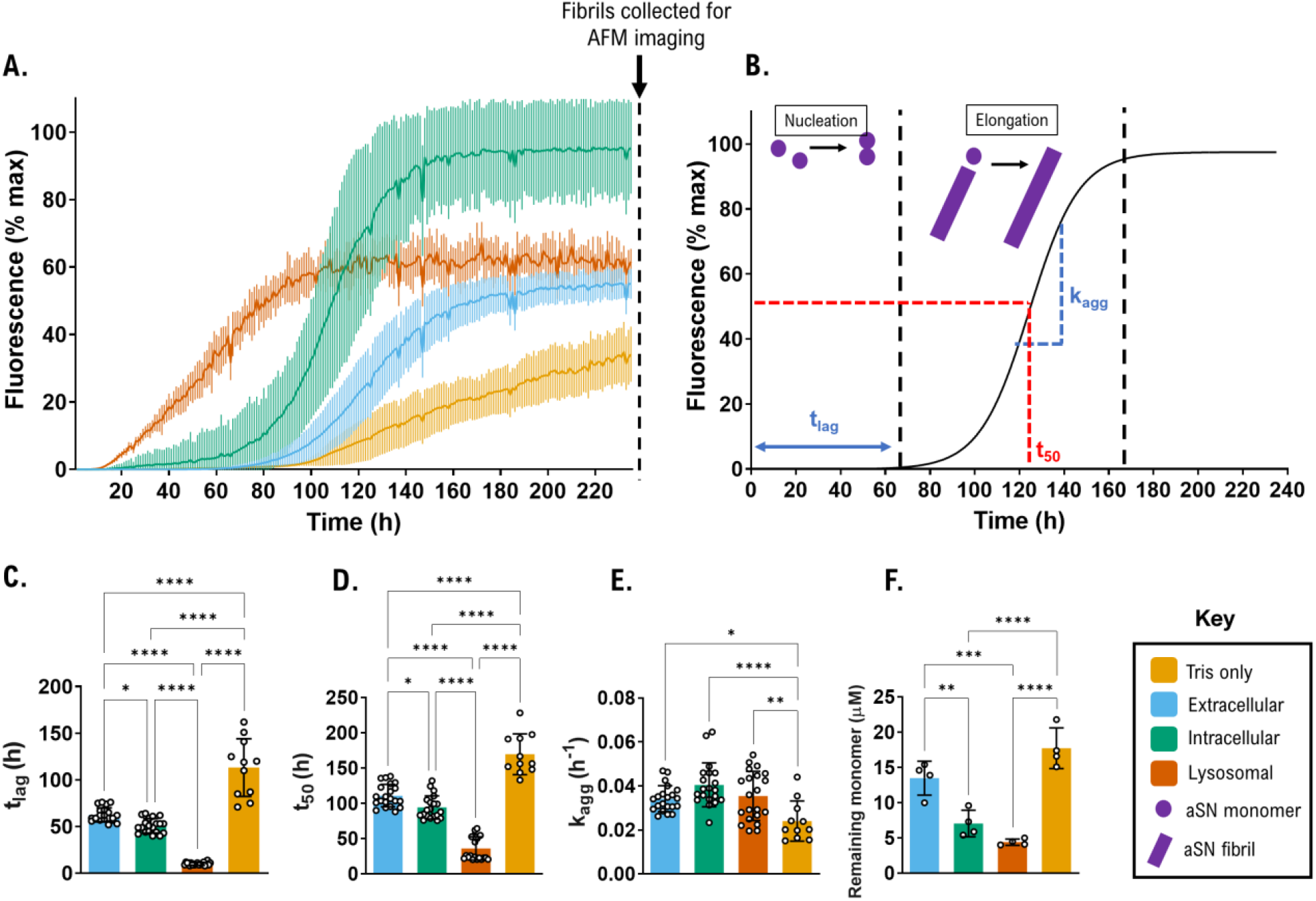
ThT-based aggregation assays reveal distinct aggregation behaviour for aSyn when equilibrated in different physiological solution conditions. **(A)** Aggregation kinetics of aSyn in Tris-only (yellow), extracellular (blue), intracellular (green) and lysosomal (orange) solution conditions were measured using ThT fluorescence intensity and plotted as % of maximum fluorescence at 480 nm. Trace shows average and standard deviation of up to 9 technical replicates. Biological replicate 1 shown (see Supplementary Figure 2 for all biological replicates); **(B)** The aggregation phases of nucleation (lag time) and elongation (slope of curve) are shown schematically. **(C-F)** Lag time (t_*lag*_), time to reach 50% of maximum aggregation (t_*50*_) and slope (k_*agg*_) were calculated and significance testing was performed by a one-way ANOVA with Tukey’s multiple comparisons post-hoc test. The upper and lower 95% confidence interval is shown and p-value significance of differences between cellular conditions are indicated (p < 0.05*, p < 0.01**, p < 0.001***, p < 0.0001****). The remaining monomer concentration was determined using SEC-HPLC by injecting 25 µL of soluble sample from each well in the ThT assay and calculating the area of the aSyn monomer peak in relation to a standard curve of known aSyn monomer concentrations. Remaining monomer concentrations were measured from the area under the peak and calculated using a standard curve of known concentrations. Data shown in C-E correspond to n=21 for the extracellular, intracellular and lysosomal conditions, and n=11 for Tris-only. Data in F correspond to n=4.

As ThT-based fluorescence intensity can change due to presence of different fibril polymorphs and solution conditions^41–43^, we confirmed the extent of aggregation by quantifying the remaining monomer concentration at the end of the assays (Figure 1F). Thus, the aggregation propensity can be described as the reciprocal of the remaining monomer. The order of aggregation propensity from highest to lowest was: Lysosomal > Intracellular > Extracellular > Tris-only. Importantly, the cellular and extracellular compartment conditions all had a higher aggregation propensity than the Tris-only condition, showing that when deprived of biological salts, Tris-only is not physiologically relevant despite being at a physiological pH of 7.4. This may be particularly significant for drug discovery efforts which often use aSyn protein in buffers without a full complement of dissolved physiological salts corresponding to the relevant physiological compartment. We also note that aggregation propensity correlates with pH — the lower the pH, the greater the aggregation propensity.

The lysosomal condition corresponds to the fastest aSyn aggregation rate, as has been previously shown^44^. The different aggregation kinetics and propensities that we observed logically provoke the question as to whether they also result in different aSyn fibril polymorphs, thus we next imaged the fibrils in each case.

### Different physiological conditions result in five distinct fibril polymorphs

Next, we examined the fibrils formed under each condition to identify any resulting morphological variations. As previous studies have shown, the morphology of aSyn fibrils are highly sensitive to solution conditions such as pH and ionic composition^45,46^. The properties of the different polymorphs such as toxicity and seeding potency may differ^24^. The AFM analysis showed that all conditions had a percentage of the total population as non-periodic, or rod fibrils, that we termed polymorph p1 (Figure 2). Tris-only fibrils were predominantly comprised of twisted polymorphs p2 (Figure 2A and 2E). These were divided into two sub-polymorphs p2a and p2b, as they both had a long periodicity of ∼400 nm, but had different heights, with polymorph p2b (12-17 nm) having approximately double the height of polymorph p2a (7-10 nm). This can be rationalised by two protofibrils (p2a) associating to form a mature fibril (p2b)^47^. Polymorph p2b formed 60% of the total fibril population, with the lower height p2a a further 24% and the rod polymorph p1 making up only 16% (Figure 2E). The extracellular conditions created a single population of protofibrils containing the periodic polymorph p3a, which was more tightly twisted than the Tris-only fibrils, with a short periodicity of ∼100 nm (Figure 2B). None of the protofibrils were found with heights more than 8 nm, suggesting they had not laterally associated or twisted together under extracellular conditions. The lysosomal condition created two periodic fibril populations: (i) polymorph p3a; indistinguishable from the extracellular condition and (ii) polymorph p3b; a mature fibril of the same periodicity but double the height of protofibril p3a (Figure 2D). In both, the extracellular and the lysosomal conditions, most of the fibril populations were of p3a (7-9 nm), comprising 80% and 59%, respectively (Figure 2F and 2H). The intracellular fibril population was more diverse and included polymorphs found across the three other conditions, with the majority (46%) of the fibrils being p2b (Figure 2C and 2G). Figure 3 shows a summary of the different twisted polymorph populations formed across the conditions tested.

**Figure 2:**
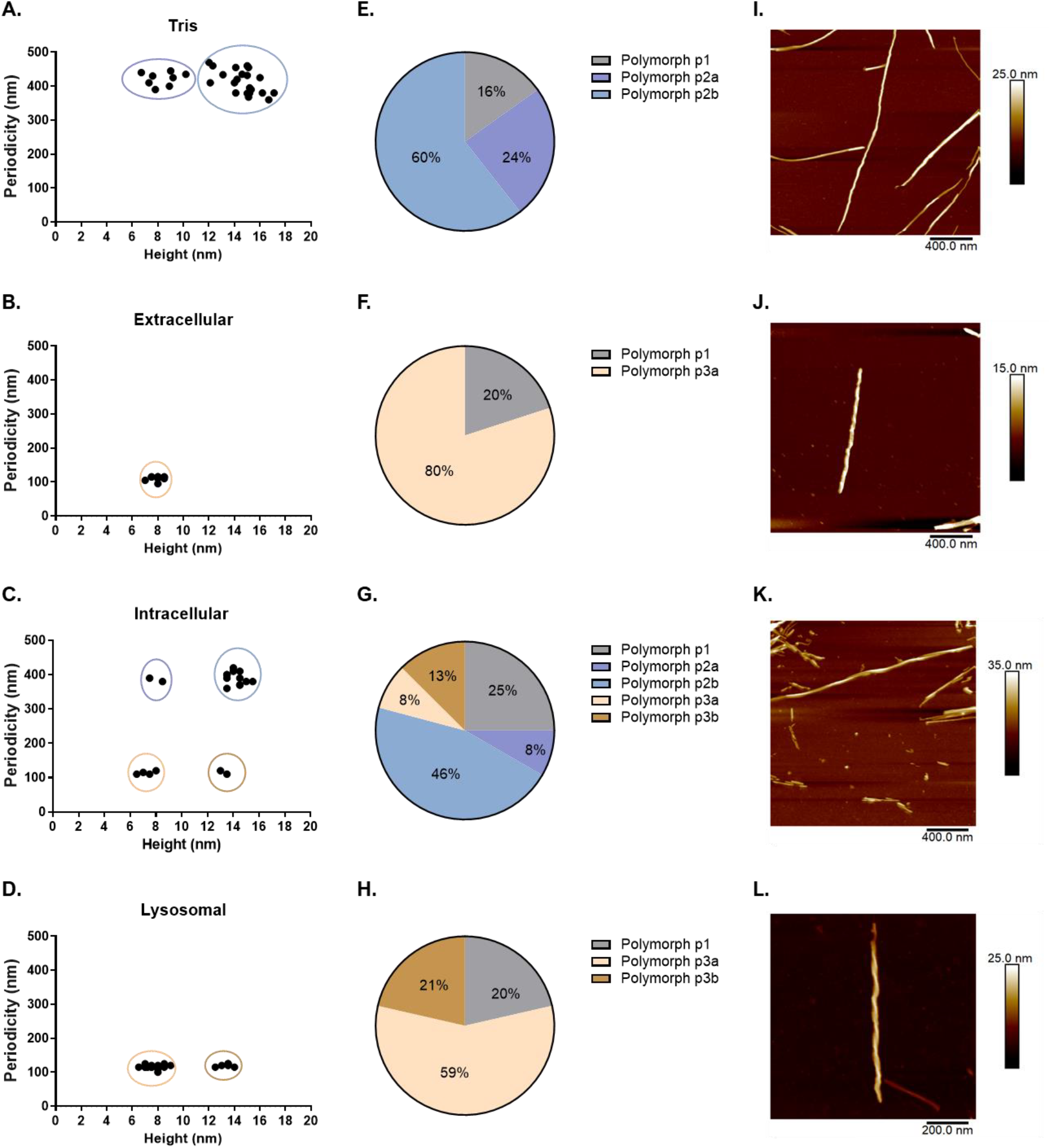
AFM analysis on the aSyn fibrils formed from each condition reveals distinct polymorphs. Atomic force microscopy was performed on the fibrils developed in each cellular, extracellular and Tris-only condition. **(A)-(D)** show plots of the periodicity against the height in nm for each condition. As a guide to the eye, groups of distinct polymorphs are highlighted by a colour ellipse. Non-periodic fibrils (or rods) are not depicted. **(E)-(F)** show pie charts representing the abundance of each polymorph population. Polymorph p1 (grey) represents fibril rods, while polymorphs p2-p3 are twisted fibrils of varying periodicities and heights, with colours matching the cluster circles in **(A)**-**(D)**. (I)-(L) show representative AFM images of the main fibril polymorph in each condition. Note scale bars and height colour bars are not identical.

**Figure 3:**
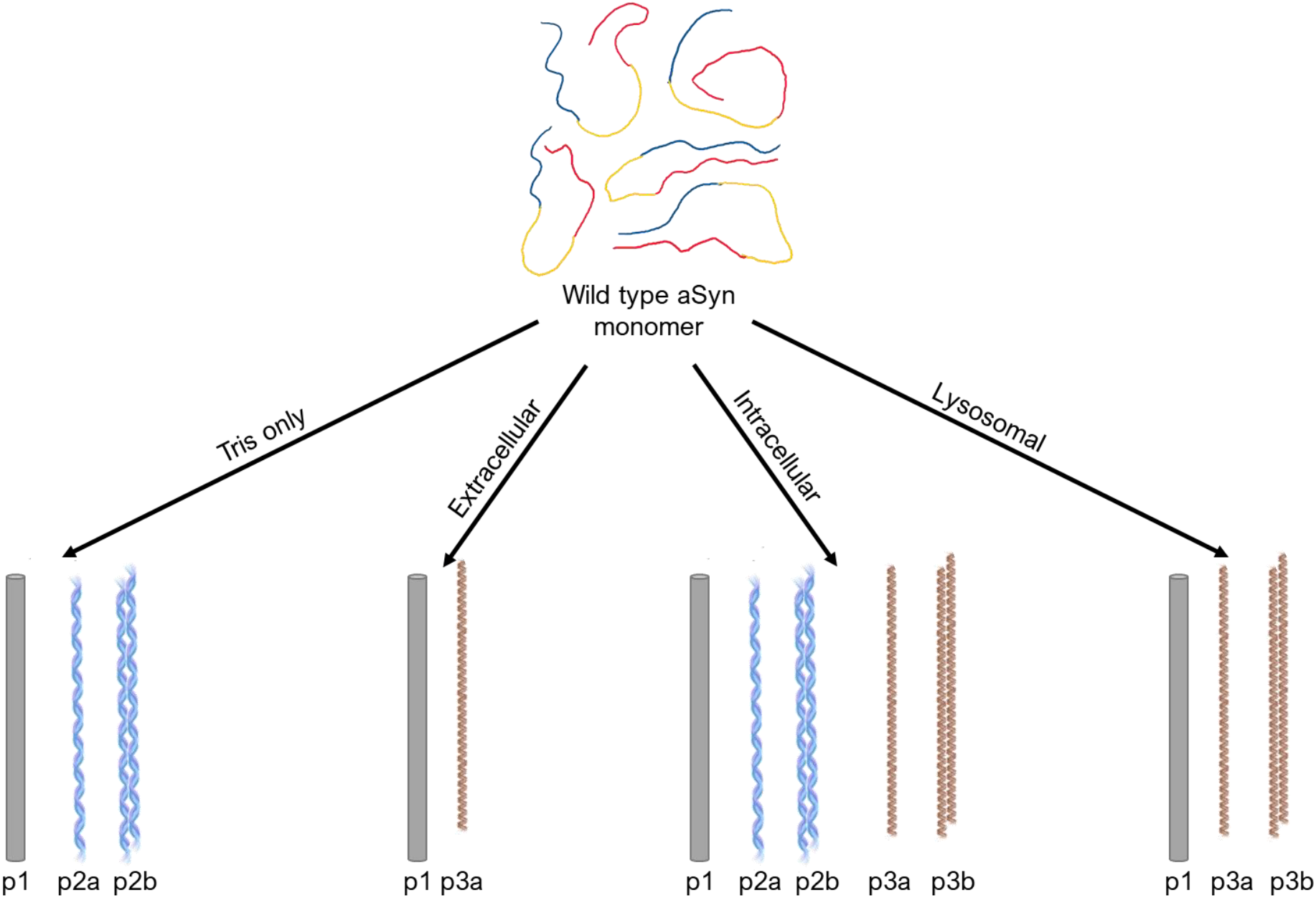
Schematic of the twisted fibril polymorphs formed from wild type aSyn for each condition. Tris-only conditions form less periodic twisted polymorphs, p2a and p2b (blue); Extracellular and lysosomal conditions form highly twisted fibrils, p3a and p3b (brown); Intracellular conditions form a mixture of the p2 and p3 polymorphs.

The AFM analysis shows that the cellular, extracellular and Tris-only conditions cause aSyn to form fibrils with five distinct morphologies, p1, p2a, p2b, p3a and p3b. We next sought to identify whether this stems from different aSyn monomer conformations under the different solution conditions.

### Monomer conformations vary with solution conditions

We hypothesised that the conformational ensemble and local structure of the aSyn monomer could be affecting the aggregation kinetics (as shown in our previous paper^21^) and the resulting fibril morphologies. To measure the local (i.e. sub-molecular) structural and conformational dynamics of the monomer in the different environmental conditions, we employed HDX-MS on the millisecond timescale. Protein conformational dynamics exquisitely influence the exchange of amide hydrogens in the polypeptide backbone, which can be sensitively measured by HDX-MS. The sub-second kinetics are essential to generate data on weakly-stable and intrinsically disordered protein monomers, such as aSyn, under physiological conditions – in particular at higher pH found in extracellular and intracellular environments^48^. Thanks to our prototype instrument^49^, we were able to capture the exchange kinetics of aSyn monomer from 50 ms, compared to a conventional lower limit of 30 s for standard commercially available HDX systems. We coupled HDX-MS with ‘soft fragmentation’ by ETD^50^ in order to further increase the structural resolution of the data, with 21% of aSyn resolved at the single amino acid level (Supplementary Figure 3). Thus, aSyn conformational perturbations can be highly localised to regions of the protein that are involved in specific processes, in this case, aggregation. These HDX-MS data measure the ensemble average of monomeric aSyn conformers formed under the physiological conditions.

Intrinsic amide hydrogen/deuterium-exchange (HDX) varies with pH and ionic strength, which must be corrected for in order to measure only the HDX differences that result from the structural dynamics of the aSyn protein. We first used the unstructured peptide bradykinin to empirically calibrate the chemical exchange rate in each solution condition^50–52^ (see *Methods*). This resulted in a set of correction factors that permit the normalisation of experimental data in each solution condition to a common scale (Supplementary Figure 4). Therefore, we were able to robustly determine which the significant conformational changes were in monomeric aSyn between the different chemical environments. Briefly, we used the hybrid significance testing method^53^, combining the results of a Welch’s t-test and determining a global significance threshold corresponding to the experimental error, to identify significant differences between the conditions for the deuterium uptake per labelling timepoint and per amino acid (see *Methods* and Seetaloo et al^50^).

Figure 4 shows the HDX-MS results as a heatmap showing only the significant differences in uptake at each experimental timepoint, from 50 ms to 30 s, in a pairwise manner between the conditions. Part of the C-terminus is significantly protected in the extracellular state, compared to the intracellular and lysosomal states (blue residues in Figure 4B-C). Conversely, the N-terminus and NAC residues 2-4, 10-17, 34-38 and 53-94 are deprotected (red residues in Figure 4B-C). A similar pattern is seen for the extracellular/intracellular vs lysosomal differential across residues 1-112, with the remainder of the C-terminal sequence showing a slightly different pattern of uptake difference. On the other hand, the comparisons of Tris-only versus all the physiological states (Figure 4E and S5) show protection against HDX throughout, with highest protection conferred to the C-terminus. The differential HDX-MS analysis confirms that the aSyn monomer varies in conformational ensemble across the physiological and Tris-only conditions studied here and localises the ensemble averaged conformational changes.

**Figure 4:**
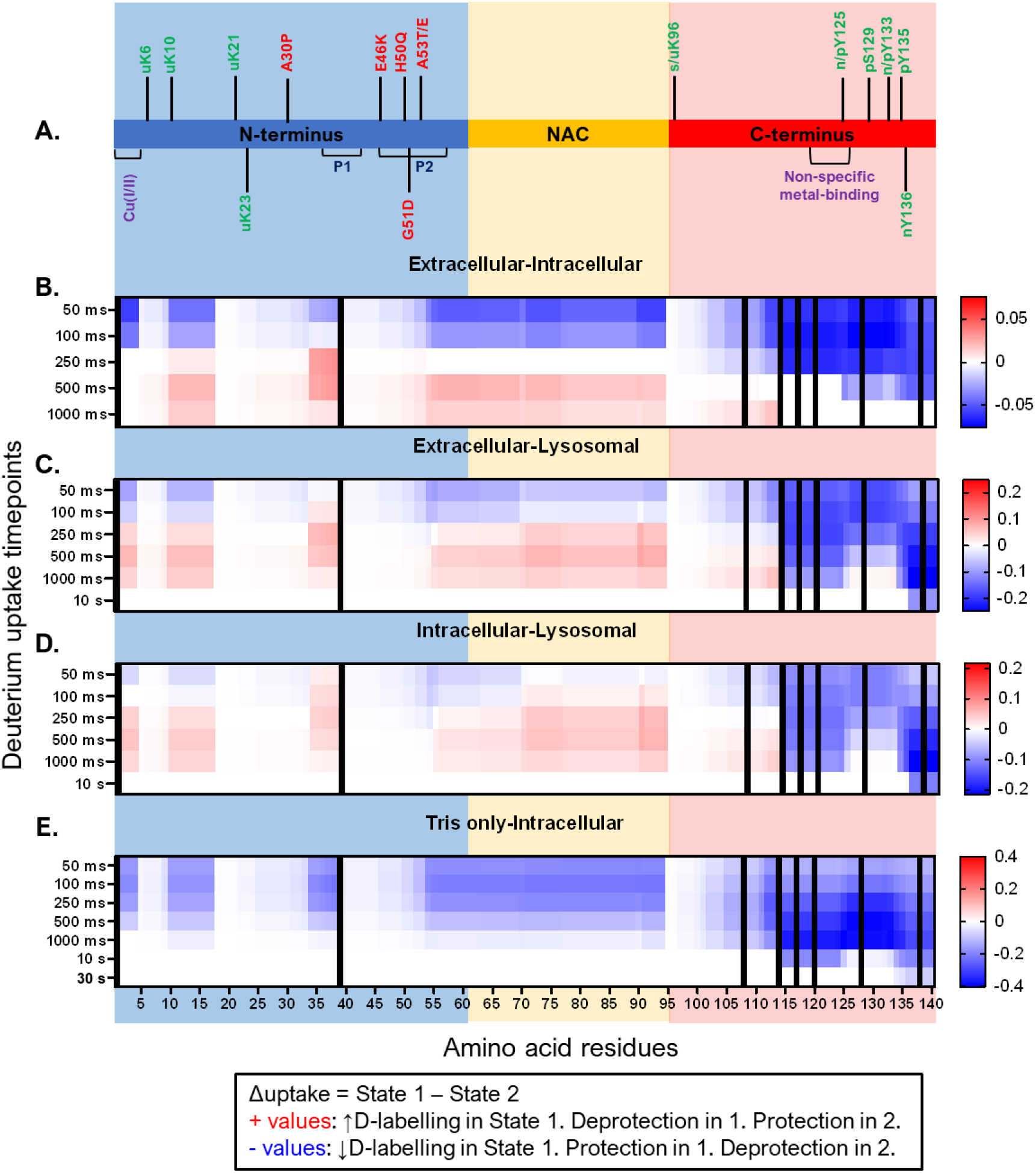
HDX-MS reveals localised differences in conformations of monomeric aSyn across all the conditions. **(A)** Schematic of aSyn monomer with important features and domains shown. **(B-E)** Heatmap showing significant differences (non-white) in deuterium uptake per timepoint during an on-exchange reaction between STATE 1 – STATE 2 (title of each plot). Hybrid significance testing with Welch’s t-test p-value of 0.05 and global significance threshold of 0.36 Da calculated. Data for three biological replicates shown. Data are resolved to the amino acid level, down to single residues in certain regions. Positive values are in red and represent increased uptake in STATE 1, whereas negative values are in blue and represent increased uptake in STATE 2. Increased uptake indicates more solvent exposure and/or less participation in stable hydrogen-bonding networks. Tris-only comparisons with extracellular and lysosomal similar to E, not shown here but in Supplementary Figure 4.

### Exposure of C-terminus residues 115-135 are key for nucleation

We then sought to correlate the localised structural perturbations in the monomeric aSyn with the nucleation and elongation phases of the aggregation kinetics. We aimed to determine if there were certain structural motifs or regions in the aSyn monomer whose protection or deprotection to HDX reveal a contribution to each aggregation phase.

We performed a Pearson correlation analysis at each amino acid in aSyn, with a 99% confidence limit, between the nucleation lag time (t_*lag*_) from ThT-based assays (Figure 1) and the observed rate constant (k_obs_) of hydrogen-exchange (Figure 5A-B; Supplementary Figure 2). Table S1 shows the Pearson correlation coefficients R at each amino acid. C-terminus residues 115-135 are very strongly negatively correlated with t_lag_ (R < -0.9), while the rest of the C-terminus is strongly negatively correlated (−0.9 < R < -0.7), albeit to a lesser extent (Figure 5B). Similarly, certain localised regions of the N-terminus (10-33 and 40-60) and the NAC region (61-69) are also strongly negatively correlated (−0.9 < R < -0.7), but to a lesser extent. Therefore, aSyn conformations, where the above-mentioned residues are exposed and/or their hydrogen-bonding networks are destabilised, are found to nucleate more rapidly.

**Figure 5:**
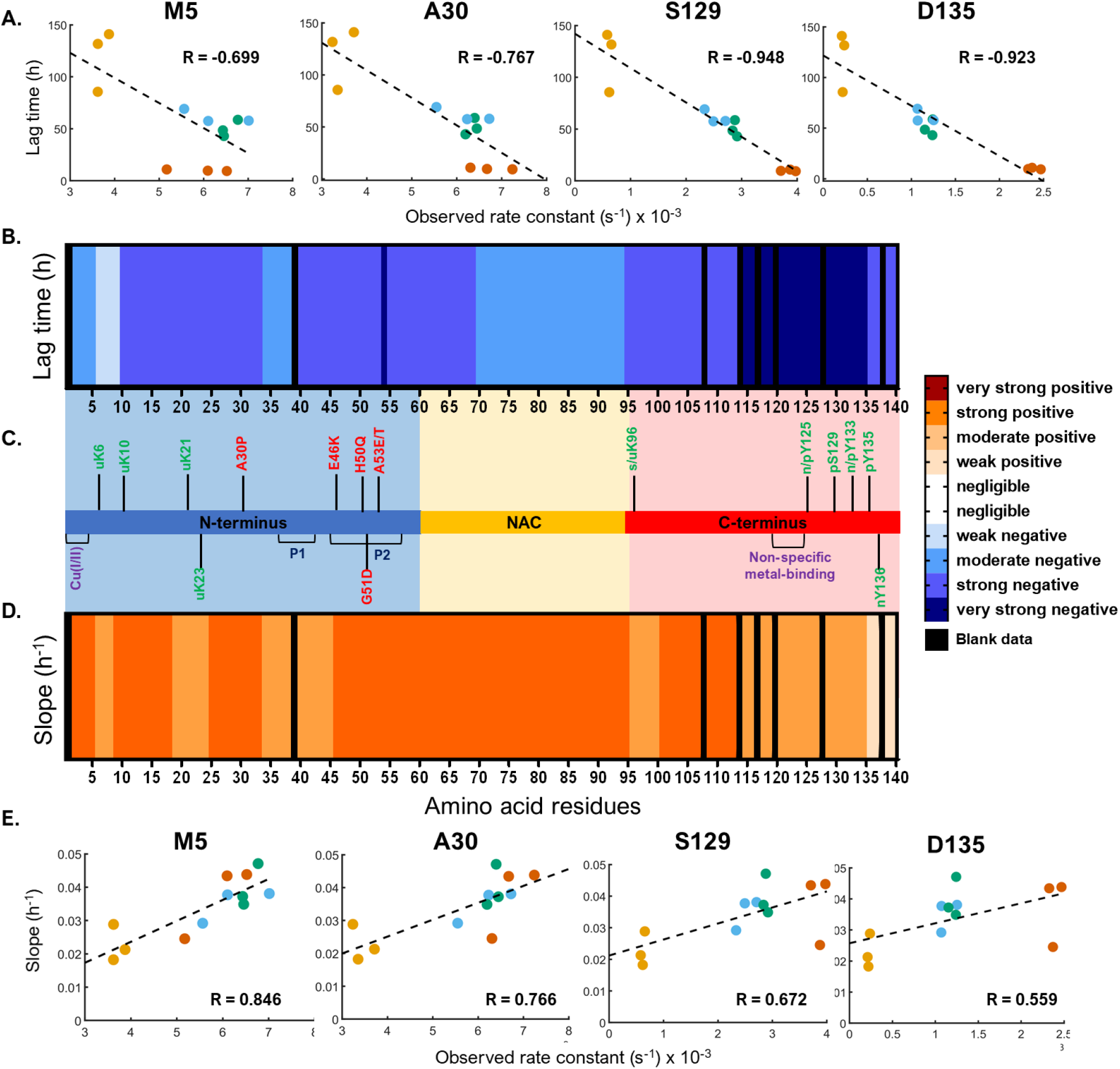
Correlation analysis reveals regions of the protein important to the nucleation and elongation phases of the aggregation kinetics. **(A)** Correlation plots of lag time against k_obs_ for selected amino acid residues – M5 (copper-binding site), A30 (familial mutation site), S129 and D135 (post-translational modification sites); **(B)** Heatmap of the Pearson correlation coefficients (R) between the lag time and k_obs_; **(C)** Schematic of the aSyn sequence showing the three domains (N-terminus in blue, NAC in yellow, C-terminus in red), sites of selected familial mutations (red), metal-binding (purple) and post-translational modifications (green); **(D)** Heatmap of the Pearson correlation coefficients (R) between the aggregation slope and k_obs_; **(E)** Correlation plots of aggregation slope against k_obs_ for selected amino acid residues (same as A). Colour bar legend shown with following categories of R: negligible: 0-0.3, weak: 0.3-0.5, moderate: 0.5-0.7, strong: 0.7-0.9, very strong: 0.9-1. Black regions represent unavailable data.

### Exposure of N-terminus and NAC regions drives fibril elongation

We next correlated the rate of fibril growth, defined by the slope of the exponential phase (k_*agg*_) from the ThT-based assays with the observed rate constant (k_*obs*_) as before. We have performed a Pearson correlation analysis, as above, and show a heatmap of the correlation coefficients R from the k_*obs*_-k_*agg*_ correlation along the protein sequence (Figure 5D-E). A strong positive correlation (0.7 < R < 0.9) can be seen between the k_*obs*_ and k_*agg*_ throughout the entire NAC region (residues 61-95) and for N-terminus residues 2-5, 10-17, 25-33, and 46-60. The C-terminal domain residues 101-113 also showed strong positive correlation coefficients (0.7 < R < 0.9) compared to the rest of the protein sequence. This means that the more exposed or less involved in hydrogen-bonding these residues are, the higher the elongation rate, implying faster fibril growth. Interestingly, C-terminal residues 115-135 that previously proved to be critical for the nucleation phase, are only moderately influential for the process of fibril elongation. Thus, monomeric conformations where the NAC region, together with the above-mentioned sites, is exposed to solvent water and/or has a destabilised H-bonding network, are found to accelerate fibril elongation.

## DISCUSSION

Being intrinsically-disordered, aSyn monomer occupies a broad but shallow conformational space as it adopts a wide range of conformations stabilised by long-range intramolecular electrostatic and hydrophobic interactions^7,8,54^. Consequently, it is challenging to define the variously meta-stable conformers using integrated structural biology tools^20,26^. Disruption of these intramolecular interactions can be caused by mutations^27–32^, local environmental changes, such as during pre-synaptic calcium signalling^21^ or by post-translational modifications^33,55–57^. This in turn can trigger misfolding and initiate aggregation^11^. This underlines the crucial importance that these physiological and patho-physiological functionally distinct conformers of aSyn are characterised and, moreover, at a structural resolution sufficient to correlate structure with functional attributes.

As aSyn experiences a plethora of chemical environments and binding partners at the presynapse, we hypothesise that its monomeric form will exhibit changes in conformation under different microenvironments. Therefore, in the current study, we have focused on whether the aSyn monomer conformation is altered under various conditions at the presynapse. We sought to mimic physiologically relevant cellular and extracellular environments (extracellular, intracellular, and lysosomal) and whether/how these conformational changes lead to distinct aggregation kinetics and fibril morphologies.

In the current study, we show that the four solution conditions under investigation: Tris-only (control), extracellular, intracellular, and lysosomal states, aggregated at different nucleation times and elongation rates, determined by ThT-based aggregation assays. Analysis of resulting fibrils by AFM revealed an assortment of fibril polymorphs across the four environments, indicating a link between the net-charge in the C-terminal domain and fibril twist. More specifically, we found that a net charge reduction at the C-terminus led to the formation of fibril polymorphs with increased periodicity. Subsequently, we were able to identify subtle perturbations in the conformational ensembles of the wild-type aSyn monomer in the physiologically relevant solution conditions at high structural resolution using millisecond HDX-MS coupled with ETD. Ultimately, we were able to determine correlations between the monomer conformers and specific stages of aggregation: nucleation and elongation.

We correlated the different cellular aggregation profiles from the ThT-based assays with the k_obs_ from HDX-MS and discovered that the C-terminal residues 115-135 crucially influenced the nucleation of the fibrils, as shown by the very strong correlation coefficients spanning this region. Previous studies have shown that the truncation or charge neutralisation of the C-terminus increases the rate of aggregation. As we saw, the k_*obs*_ for HDX at the C-terminus strongly negatively correlated with t_*lag*_, indicating that the more exposed it is or the less involved in stable H-bonding network, the faster the nucleation. When the C-terminus is truncated, which mimics a fully exposed/unstructured conformation, this leads to increased aggregation, which is also observed when the C-terminus charge is neutralised^58,59^. Upon charge reduction at the C-terminus (possibly via calcium-binding or lowering of pH), a drop in the long-range interactions and electrostatic repulsions may lead to increased exposure or destabilised H-bonding participation, resulting in faster nucleation. On the other hand, some residues (2-5, 10-18, 25-33, 46-60, 61-95 and 100-113) were found to promote fibril growth when deprotected against HDX (deep orange in Figure 6D). Perhaps unsurprisingly, the entire NAC was found to be highly important in the process of fibril growth, in agreement with previous deletion and truncation studies^60^. Furthermore, recent studies have identified two motifs at the N-terminus, P1 (residues 36-42) and P2 (residues 45-57) to be critical for aggregation^61^. Our high-resolution analysis shows that exposure of motif P2 drives both nucleation and fibril growth processes of aggregation, and to a higher extent than that of motif P1. The HDX-MS differential analysis revealed that P1 and P2 were more exposed in the extracellular state compared to both the intracellular and lysosomal states (red residues in Figure 4B-C).

Interestingly, we observed that fibrils formed under extracellular and lysosomal conditions led to the same more tightly twisted fibrils, polymorph p3^62^. The only significant difference between the aggregates formed under these two conditions was the propensity of the lysosomal buffer to drive assembly of p3a protofibrils into p3b mature fibrils. In common, both conditions lead to a net charge reduction at the C-terminus, either by calcium-binding^63^ or neutralisation of certain acidic residues at the lower pH^64^, respectively, which would disrupt the long-range electrostatic and hydrophobic interactions that stabilise the monomer in solution^21^. It is likely that a change in the protofibril structure and/or charge halts the formation of mature fibrils by affecting their association. CryoEM studies have revealed the formation of a different aSyn polymorph upon the E46K point mutation, which led the protofibrils to adopt a different fold compared to previously resolved wild-type aSyn structures^65,66^. It is possible that a different monomer conformation (lysosomal vs extracellular) could lead to different protofibril packing and reduced stability of the mature fibril. This suggests that the same aggregation pathway may be followed to generate the p3a fibrils from the aSyn monomer in the extracellular and lysosomal environments, but that the mature fibrils have considerably higher stability under the lysosomal conditions. From our HDX-MS vs ThT correlative analyses, the C-terminus deprotection was also found to correlate with the nucleation phase of aggregation, agreeing with previous work^58,67,68^. Therefore, we can infer that polymorph p3 is determined by an aSyn monomeric conformation with a C-terminus with lower net charge during the nucleation phase.

The intracellular condition formed the most heterogeneous fibril populations out of the four conditions, as it had all the polymorphs of the other conditions combined. It also gives rise to the widest range of fibril elongation rates (Figure 1E). The ensemble average of structural conformers, as measured by HDX-MS, was broadly similar between intra/extra-cellular conditions, however, the intracellular environment stabilises specific sites in the N-terminal region and to a far greater degree destabilises the C-terminal region (Figure 4B). The C-terminal protection can be attributed to calcium binding^69^. The intracellular state also contains Mg^2+^, which is known to bind to aSyn^20^. It is possible that in this case, Ca^2+^ binds preferentially to the Mg^2+^, but this statement can only be confirmed if a direct comparison of the two ions is performed (e.g., Tris + Ca^2+^ vs Tris + Mg^2+^). Together, these results suggest that the intracellular state stabilises aSyn in a relatively diverse set of monomeric conformations and net charge states and that these aggregate into a heterogeneous mixture of fibrils, which could be associated with different biophysical properties, levels of toxicity and disease-relevance.

It is important to note that while this study presents correlations between local structural dynamics and aggregation in wild-type aSyn, there are a wide variety of familial mutations, post-translational modifications, and even different physiological buffers – all of which have the potential to change those site-specific correlations. For example, in the case of mutation H50Q, where a basic residue is swapped for an amidic one, the electrostatics are changed with removal of a formal charge, which may impact on the specific chemistry involved in nucleation/elongation processes, and the observed rate constant would decrease by 4.2x based on the intrinsic rates documented by Bai et al^70^. This would likely affect the correlation at this residue and any other structurally connected sites elsewhere in the protein. Thus, each aSyn variant and the chemical environment should be considered non-trivial to extrapolate and each deserves assessment.

In the present study, we found that deprotection in the centre of the C-terminal domain was found to be significantly correlated with the nucleation phase of the aggregation kinetics and we identified specific residues that influenced fibril growth. We also discovered that the morphology of certain fibril polymorphs was determined as early as during monomer nucleation. We anticipate that in the future, the tools and generally applicable approach that we present here will be able to make further important structure-function correlations for other physiological conditions and proteoforms of aSyn.

## METHODS

### Materials

All media and reagents were purchased from Sigma-Aldrich (UK) and were of analytical grade unless otherwise stated. Deuterium oxide (99.9% D_2_O) was purchased from Goss Scientific (catalogue number: DLM-4). E. coli BL21STAR (DE3) cells were purchased from Invitrogen (USA). Peptide P1 was synthesised using the method described in Phillips et al^71^. Details about the expression and purification of wild-type alpha-synuclein have been described previously^21,72^. aSyn refers to the wild-type variant of the protein in this paper. Three biological replicates were produced for the use in all experiments.

### Sample preparation

Four buffer conditions were used in this study: Tris-only, extracellular, intracellular and lysosomal^15^ (Table 1). aSyn samples from three different purification batches were removed from -80°C storage (15-25 µM stocks in Tris-only equilibrium buffer). For the Tris-only sample, the protein concentration was adjusted to 5 µM with Tris-only equilibrium buffer. For the extracellular sample, the salts were directly diluted into the 5 µM protein sample. For the intracellular and lysosomal samples, the protein was buffer exchanged into the matching equilibrium buffer for six cycles using 3K MWCO regenerated cellulose Amicon ultra centrifugal filters (Millipore, USA) and made to 5 µM.

### Hydrogen-deuterium exchange mass spectrometry of alpha-synuclein samples

For labelling times ranging between 50 ms and 5 min, hydrogen-deuterium exchange (HDX) was performed using a fully-automated, millisecond HDX labelling and online quench-flow instrument, ms2min^49^ (Applied Photophysics, UK), connected to an HDX manager (Waters, USA). For each cellular condition and three biological replicates, aSN samples in the equilibrium buffer were delivered into the labelling mixer and diluted 20-fold with labelling buffer at 20°C, initiating HDX. The duration of the HDX labelling depended on the mixing loops of varying length in the sample chamber of the ms2min and the velocity of the carrier buffer. The protein was labelled for a range of times from 50 ms to 5 min. Immediately post-labelling, the labelled sample was mixed with quench buffer in a 1:1 ratio in the quench mixer to arrest HDX. The sample was then centred on the HPLC injection loop of the ms2min and sent to the HDX manager. For longer timepoints above 5 min, a CTC PAL sample handling robot (LEAP Technologies, USA) was used. Protein samples were digested onto an Enzymate immobilised pepsin column (Waters, USA) to form peptides. The peptides were trapped on a VanGuard 2.1 × 5 mm ACQUITY BEH C18 column (Waters, USA) for 3 minutes at 125 µL/min and separated on a 1 × 100mm ACQUITY BEH 1.7 μm C18 column (Waters, USA) with a 7-minute linear gradient of acetonitrile (5-40%) supplemented with 0.1% formic acid. Peptide samples did not require the initial peptic digestion step. The eluted peptides were analysed on a Synapt G2-Si mass spectrometer (Waters, USA). An MSonly method with a low collisional activation energy was used for peptide-only HDX and an MS/MS ETD fragmentation method was used for HDX-MS-ETD. Deuterium incorporation into the peptides and ETD fragments was measured in DynamX 3.0 (Waters, USA).

### ETD fragmentation of aSN peptides

The ETD reagent used was 4-nitrotoluene. The intensity of the ETD reagent per second, determined by the glow discharge settings, was tuned to give a signal of approximately 1e7 counts per second (make-up gas flow: 35 mL/min, discharge current 65 µA) to give efficient ETD fragmentation. Instrument settings were as follows: sampling cone 30 V, trap cell pressure 5e-2 mbar, trap wave height 0.25 V, trap wave velocity 300 m/s, transfer collision energy 8 V and transfer cell pressure 8e-3 mbar. Hydrogen-deuterium scrambling was measured using Peptide P1 under the same instrument conditions (Supplementary Figure 6).

### Data analysis

The raw data was processed, and assignments of isotopic distributions were reviewed in DynamX 3.0 (Waters, USA). The post-processing analysis was performed using HDfleX^50^. Briefly, the back-exchange-corrected data points for each peptide and ETD fragment were fitted using equation 1 in one-phase.

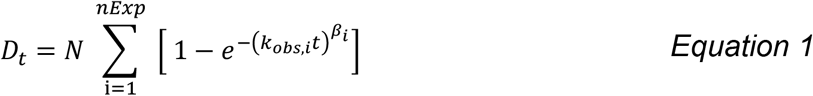

where *D*_*t*_ is the deuterium incorporation at time *t, nExp* is the number of exponential phases, *N* is the maximum number of labile hydrogens, *k*_*obs*_ is the observed exchange rate constant and *β* is a stretching factor.

As the rate of HDX is affected by pH and ionic strength, which are not controlled in this study, it is crucial to normalise the solution effects between the different conditions being compared. Here, we used an empirical approach to normalisation using the unstructured peptide bradykinin (RPPGFSPFR)^51,52^ to deconvolute the solution effects of the HDX from the protein structural changes. Due to the unstructured nature of bradykinin, all the differences in deuterium uptake seen from the different buffers can be assumed to be strictly from the changes in the chemical exchange rate effects, rather than structural effects. By using bradykinin to calibrate the chemical exchange rate, we can now clearly distinguish the structural changes between the cellular conditions (Supplementary Figure 3). The ETD fragments were combined with the peptide data using HDfleX^50^ to give the absolute uptake information across the entire protein as in Supplementary Figure 7.

### Statistical significance analysis

The hybrid significance testing method along with data flattening used here as described elsewhere^50^.

### Thioflavin-T (ThT) Binding Assay in 96-Well Plate

Thioflavin – T (ThT) kinetic assays were used to monitor the aggregation of aSyn in different cellular compartments. For sample preparation, 40 μM (final concentration) of freshly made ThT solution (Abcam, Cambridge, UK) in distilled water was added to 50 μL of 40 μM, aSyn in 20 mM Tris pH 7.4, extracellular, intracellular, and lysosomal conditions as described in Table 1.

All samples were loaded in nonbinding, clear 96-well plates (Greiner Bio-One GmbH, Germany) which were then sealed with a SILVERseal aluminium microplate sealer (Grenier Bio-One GmbH). Fluorescence measurements were taken with FLUOstar Omega microplate reader (BMG LABTECH GmbH, Ortenbery, Germany). Excitation was set at 440 nm and the ThT fluorescence intensity was measured at 480 nm emission with a 1300 gain setting. The plates were incubated with double orbital shaking for 300 s before the readings (every 60 min) at 300 rpm. Three repeats were performed with 6 replicates per condition. Each repeat was performed with a different purification batch of aSyn (biological replicate). Data were normalised to the well with the maximum fluorescence intensity for each plate and the empirical aggregation parameters t_lag_, t_50_, k, were calculated for each condition, based on the equation:

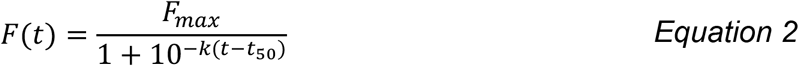

where *F* is the normalised fluorescence to the highest value recorded in the plate repeat, *F*_*max*_is the maximum fluorescence at the plateau, *k* is the slope of the exponential phase of the curve, and *t*_50_ is the time when 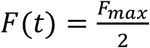.

One-way ANOVA was used to calculate statistical significance between samples using GraphPad Prism 8 (GraphPad Software, USA).

### SEC-HPLC (Size exclusion–High-performance liquid chromatography)

At the end of the ThT-based aggregation assays, the amount of remaining monomer of aSyn in each well was determined by analytical size exclusion chromatography with HPLC (SEC-HPLC). SEC analysis was performed on the Agilent 1260 Infinity HPLC system (Agilent Technologies, UK) equipped with an autosampler and a diode-array detector using a AdvanceBio 7.8 × 300mm 130 Å SEC column (Agilent Technologies, UK) in 20 mM Tris pH 7.4 at 0.8 mL/min flow-rate. 25 μL of each sample was injected onto the column and the elution profile was monitored by UV absorption at 220 and 280 nm. The area under the peak in the chromatogram of absorption at 280 nm was determined and used to calculate the monomer concentration. Monomeric aSyn samples spanning from 5 μM to 40 μM aSyn were used to determine a standard curve, to allow calculation of the protein concentration for the ThT-based aggregation assay samples based on their area under the peak.

### AFM analysis of fibril morphology

Fibrils formed at the end of ThT assays were analysed by AFM. A freshly cleaved mica surface was coated in 0.1% poly-l-lysine, washed with distilled H_2_O thrice and dried under a stream of nitrogen gas. Samples from the microplate wells were then incubated for 30 min on the mica surface. The sample was washed thrice in the buffer of choice (for example, in 20 mM Tris, pH 7.4 for the Tris condition) to remove lose fibrils. Images were acquired in fluid using tapping mode on a BioScope Resolve AFM (Bruker, USA) using ScanAsyst-Fluid+ probes. 512 lines were acquired at a scan rate of 1.5 Hz per image with a field of view of 2-5 µm and for at least ten fields of view. Images were adjusted for contrast and exported from NanoScope Analysis 8.2 software (Bruker). Measurements of fibril height and periodicity were performed by cross-sectioning across the fibril and across the fibril axis in NanoScope Analysis 8.2 software (Bruker). Statistical analysis of the height and periodicity measurements was performed in GraphPad Prism 8 (GraphPad Software, USA).

## ACKNOWLEDGEMENTS

NS is funded by a University Council Diamond Jubilee Scholarship (Exeter). JJP is supported by a UKRI Future Leaders Fellowship [Grant number: MR/T02223X/1]. GSK acknowledges funding from the Wellcome Trust (065807/Z/01/Z) (203249/Z/16/Z), the UK Medical Research Council (MRC) (MR/K02292X/1), Alzheimer Research UK (ARUK) (ARUK-PG013-14), Michael J Fox Foundation (16238) and Infinitus China Ltd. ADS and MZ acknowledge Alzheimer Research UK for travel grants. MZ acknowledges funding from Newnham College (Cambridge) and the George and Marie Vergottis Foundation (Cambridge Trust) and the British Biophysical Society (BSS) for travel grants. We thank Dr Ioanna Mela for discussions on AFM for aSyn fibril morphology. For the purpose of open access, the author has applied a CC BY public copyright licence to any Author Accepted Manuscript version arising from this submission.

## COMPETING INTERESTS

The authors declare no competing interests.

## AUTHOR CONTRIBUTIONS

NS and MZ contributed equally. NS and JJP designed the study. NS, MZ and ADS prepared proteins for experiments. NS performed HDX-MS and ETD. NS and JJP performed correlative analyses. MZ performed kinetic aggregation assays and AFM experiments. NS and MZ analysed data. NS, MZ, ADS, GSK and JJP contributed to paper writing.

## Data Availability statement

The authors declare that the data supporting the findings of this study are available in this paper and its supplementary information files. Source data are provided with this paper. All mass spectrometry .raw files will be available from the PRIDE repository [accession pending].

## Code Availability statement

This study uses in-house developed software available to download: [http://hdl.handle.net/10871/127982]^50^. Supporting information shows additional code used to calculate the Pearson correlation coefficients and to plot the correlation plots at each amino acid.

## Supporting Information

ThT, AFM and HDX mass spectrometry source data are provided in SourceData.xlsx.

## Notes

### Competing Interest Statement

The authors have declared no competing interest.

