## Supplementary Information for "Local structural dynamics of alpha-synuclein correlate with aggregation in different physiological conditions"

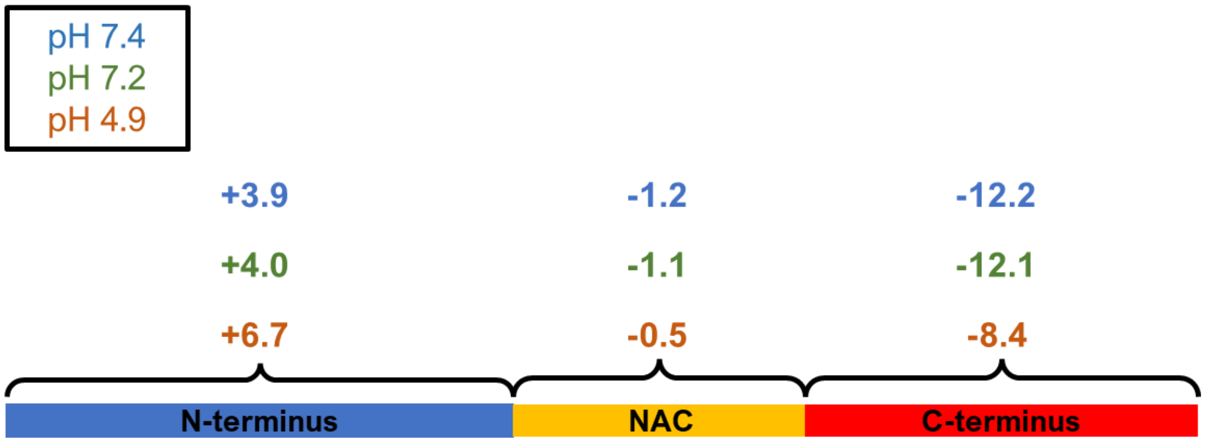

**Figure S1: Charge distribution over the alpha-synuclein sequence over pH 7.4 (blue), 7.2 (green) and 4.9 (orange).**

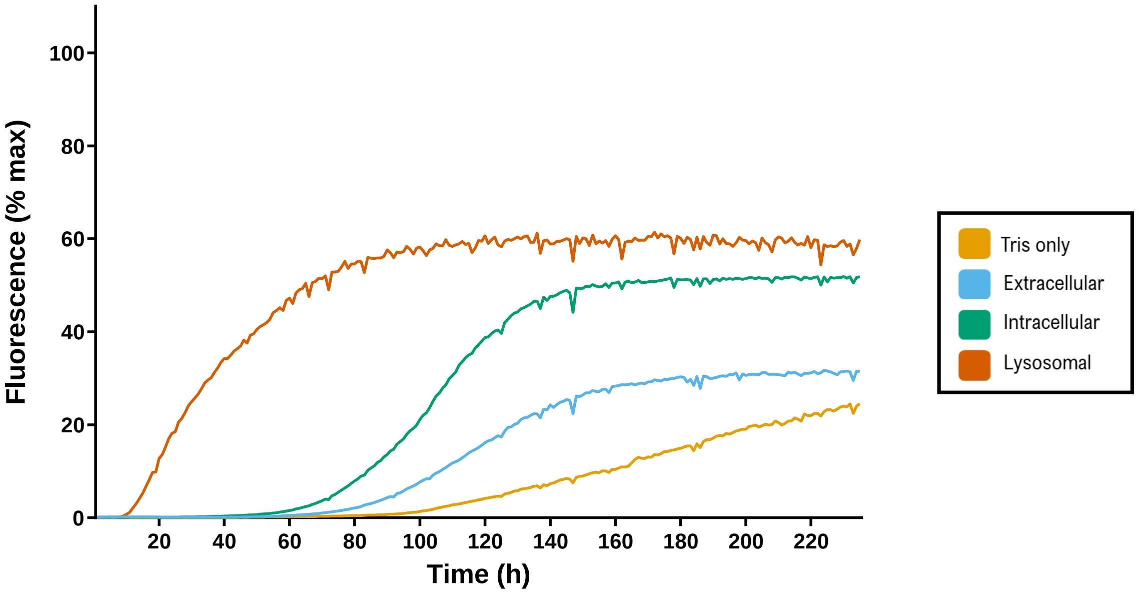

**Figure S2: ThT-based aggregation assays reveal distinct aggregation behaviour for aSyn when equilibrated in different physiological solution conditions.** Aggregation kinetics of aSyn in Tris only (yellow), extracellular (blue), intracellular (green) and lysosomal (orange) solution conditions were measured using ThT fluorescence intensity and plotted as % of maximum fluorescence at 480 nm. Trace shows the average of up to 9 technical replicates for three biological replicates; A monomer concentration of 40 μM and ThT concentration of 40 μM were used. Measurements taken every hour, with 5 min shaking before the reading. Excitation/emission at 440/480 nm, gain settings 800, 1200, 1300, 1400, 1500. Assay length 235 h. Temperature set at 37 ˚C.

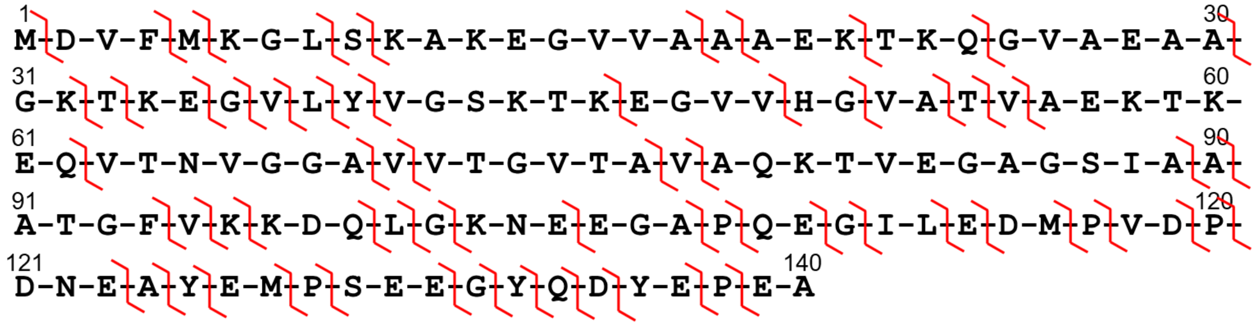

**Figure S3: Structural resolution of aSyn HDX data.** Red ticks represent the sites of either proteolytic cleavage or ETD fragmentation. The merged resolution obtained by combining peptides and c/z fragments is given by the length of the gaps between these red ticks – shorter gaps indicate higher structural resolution, with 21% of the aSyn protein at single amino acid resolution.

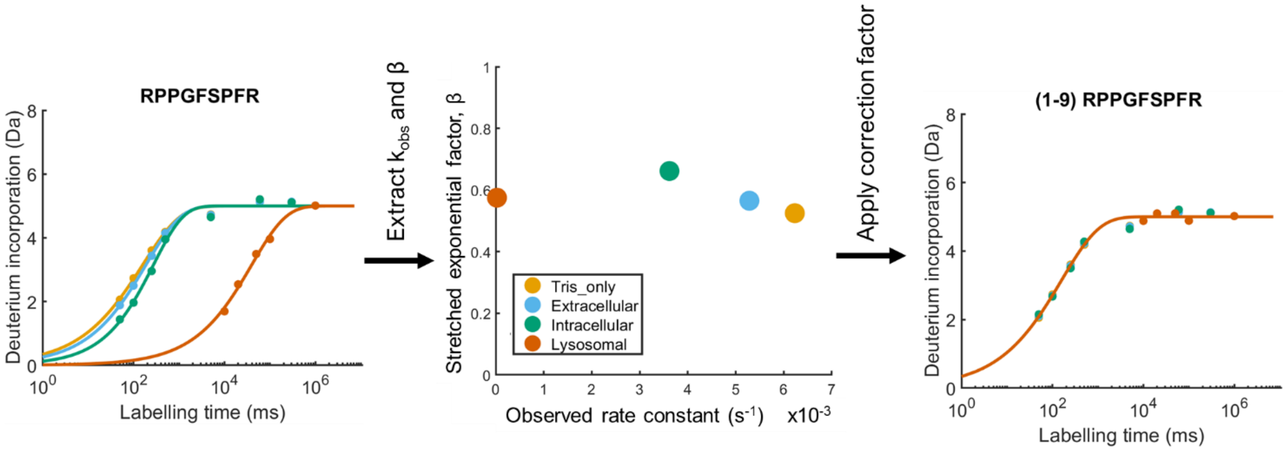

**Figure S4: Unstructured peptide bradykinin calibrates chemical exchange rate effects**; left plot: uptake curve for bradykinin in the four conditions; middle plot: 2D plot of extracted fitted parameters $k_{obs}$ and β, with Tris (yellow dot) chosen as reference state; right plot: empirical correction applied to all states. Data points are the mean of n=3 technical replicates. Note that in the right plot several data points are not visible due to overlap of all data collected at any given labelling time after the correction for the different intrinsic HDX chemistry under the different conditions.

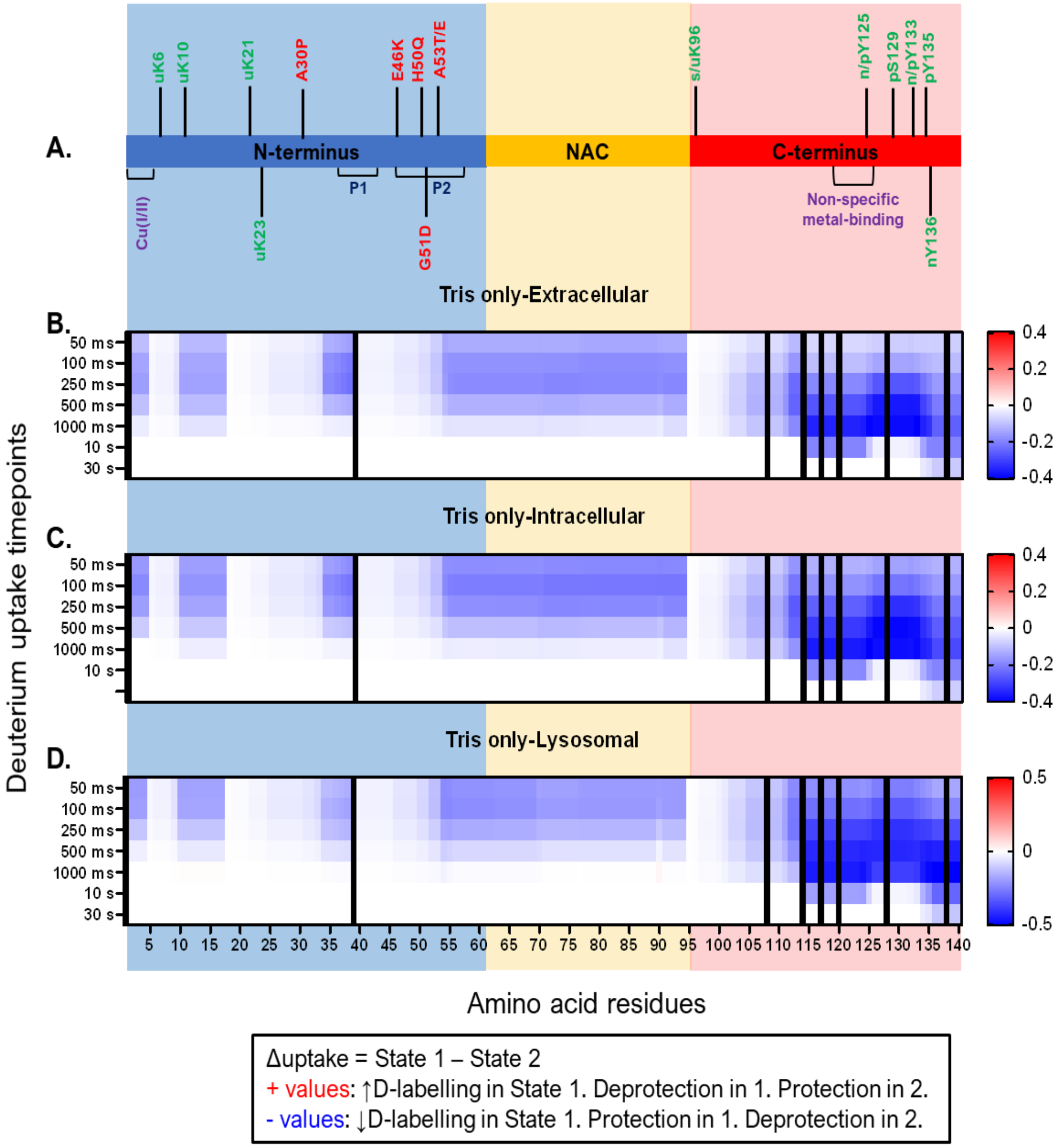

**Figure S5: HDX-MS reveals localised differences in conformations of monomeric aSyn across Tris only vs the compartment conditions.** (A) Schematic of aSyn monomer with important features and domains shown. (B-D) Heatmap showing significant differences in deuterium uptake per timepoint during an on-exchange reaction between STATE 1 – STATE 2 (title of each plot). Data are resolved to the amino acid level, down to single residues in certain regions. Positive values are in red and represent increased uptake in STATE 1, whereas negative values are in blue and represent increased uptake in STATE 2. Increased uptake indicates more solvent exposure and/or less participation in stable hydrogen-bonding networks. A possible explanation for the general protection across Tris-only could be that the salts in the cellular and extracellular buffers might be charge-shielding the N- and C-termini, such that in Tris-only, devoid of any biological salts, would adopt a more closed conformation due to increased intramolecular interactions.

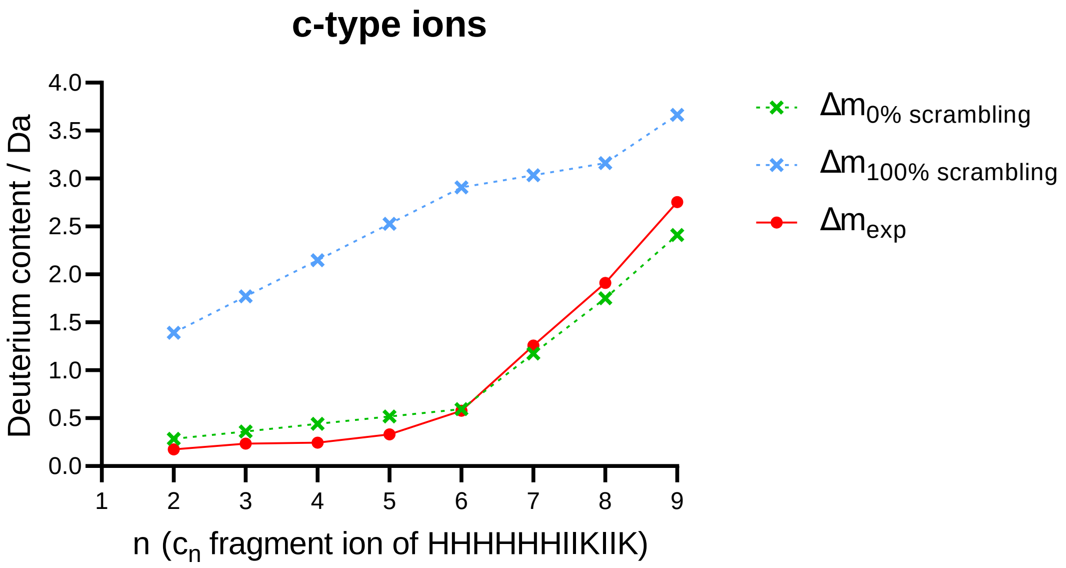

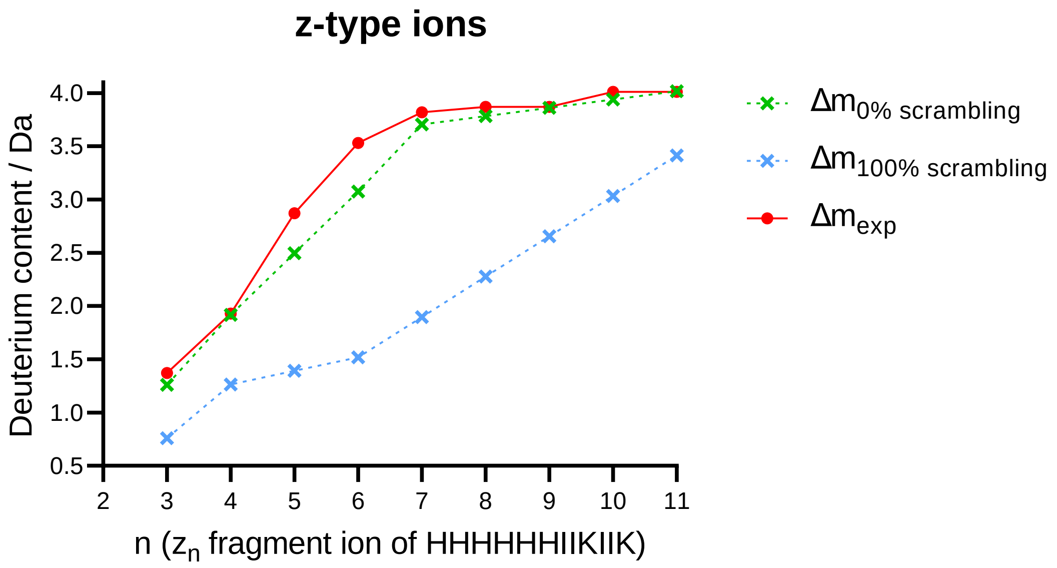

**Figure S6: Hydrogen-deuterium scrambling is not observed in the c and z fragments of peptide P1 under identical conditions as aSyn experiments.** The green and blue dotted lines on the plots represent the 0% and 100% theoretical scrambling data for this peptide respectively, and the red line shows the experimental data. From the plots, it can be inferred that the conditions chosen for ETD fragmentation do not cause H/D scrambling as the experimental data line (red) approaches the 0% scrambling line (green) for both types of fragments.

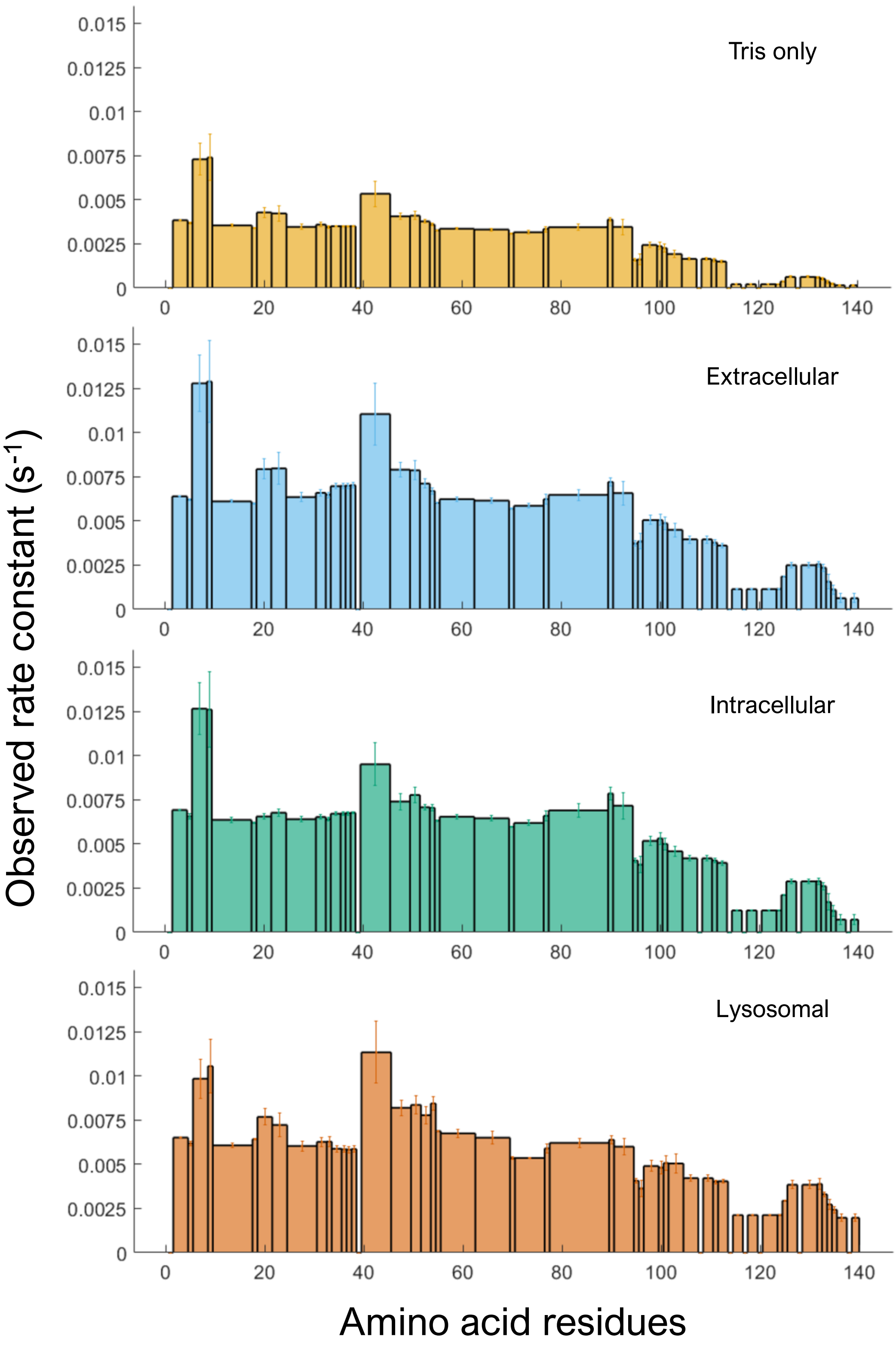

**Figure S7: HDX-MS reveals different conformations in monomeric aSyn across all the conditions.** Bars represent the observed hydrogen-exchange rate constant, $k_{obs}$, localised to the amino acids covered by the width of each bar. Error bars denote +/- 0.5 standard deviations. The N-terminus is shaded in blue, the NAC region in yellow and the C-terminus of aSyn in red. Data are for one technical replicate of three biological replicate aSyn samples. Grey boxes highlight example local features that are structurally dynamic, as evidenced by the alteration in the pattern of HDX observed rate constant between the conditions.

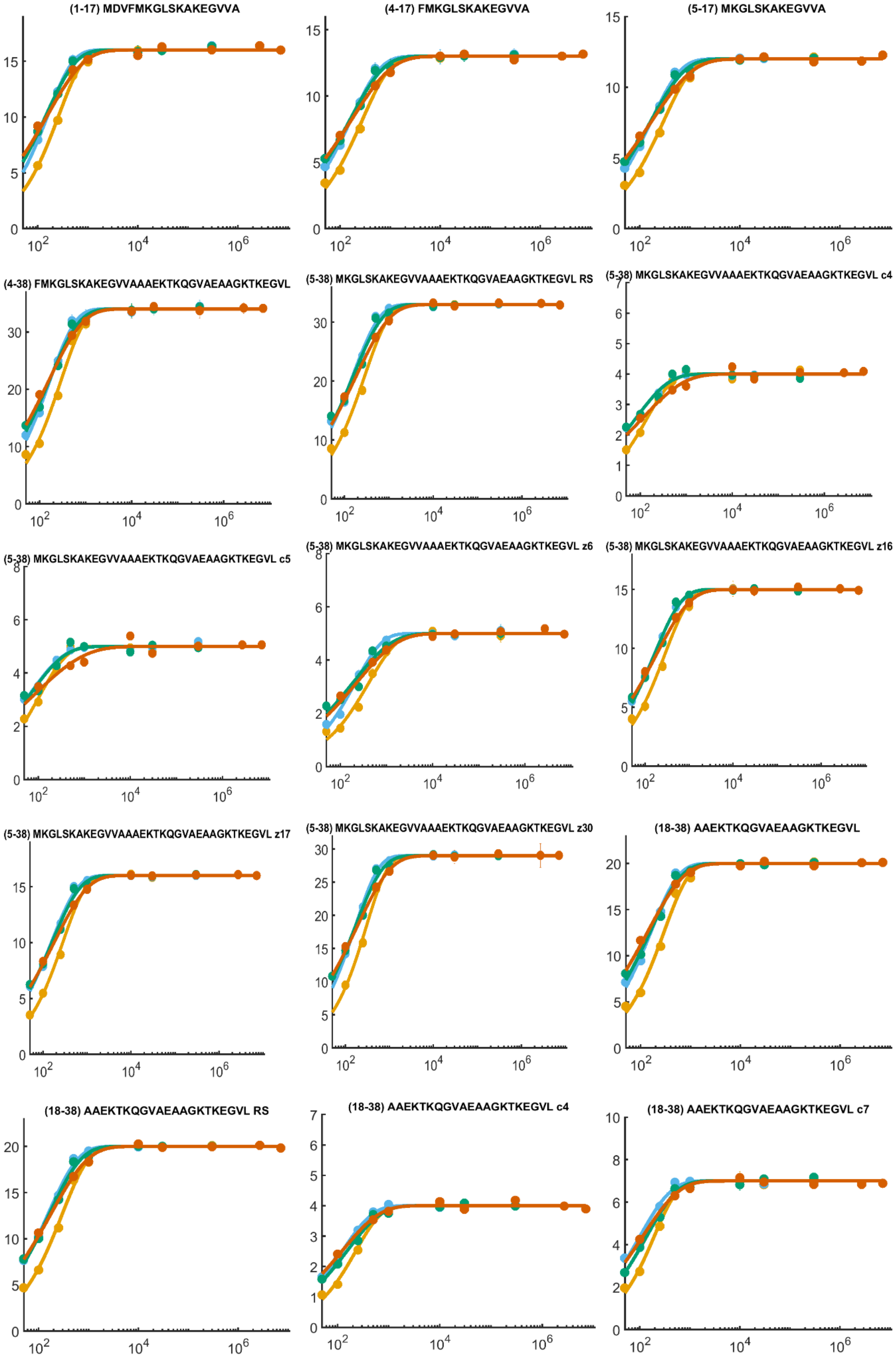
(1-17) MDVFMKGLSKAKEGVVA
(4-17) FMKGLSKAKEGVVA
(5-17) MKGLSKAKEGVVA
(4-38) FMKGLSKAKEGVVAAAEKTKQGVAEAAGKTKEGVL
(5-38) MKGLSKAKEGVVAAAEKTKQGVAEAAGKTKEGVL RS
(5-38) MKGLSKAKEGVVAAAEKTKQGVAEAAGKTKEGVL c4
(5-38) MKGLSKAKEGVVAAAEKTKQGVAEAAGKTKEGVL c5
(5-38) MKGLSKAKEGVVAAAEKTKQGVAEAAGKTKEGVL z6
(5-38) MKGLSKAKEGVVAAAEKTKQGVAEAAGKTKEGVL z16
(5-38) MKGLSKAKEGVVAAAEKTKQGVAEAAGKTKEGVL z17
(5-38) MKGLSKAKEGVVAAAEKTKQGVAEAAGKTKEGVL z30
(18-38) AAEKTKQGVAEAAGKTKEGVL
(18-38) AAEKTKQGVAEAAGKTKEGVL RS
(18-38) AAEKTKQGVAEAAGKTKEGVL c4
(18-38) AAEKTKQGVAEAAGKTKEGVL c7

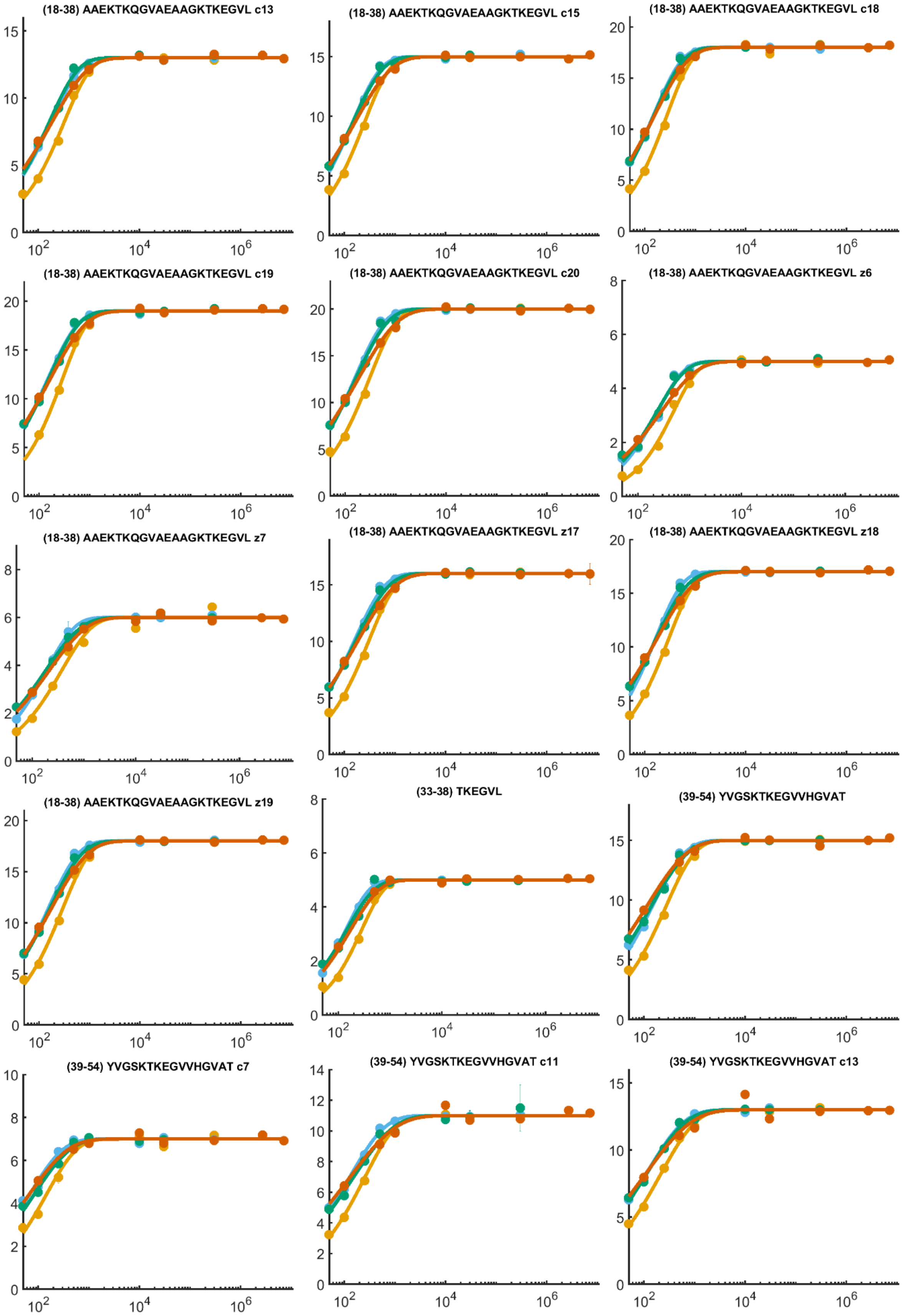
(18-38) AAEKTKQGVAEAAGKTKEGVL c13
(18-38) AAEKTKQGVAEAAGKTKEGVL c15
(18-38) AAEKTKQGVAEAAGKTKEGVL c18
(18-38) AAEKTKQGVAEAAGKTKEGVL c19
(18-38) AAEKTKQGVAEAAGKTKEGVL c20
(18-38) AAEKTKQGVAEAAGKTKEGVL z6
(18-38) AAEKTKQGVAEAAGKTKEGVL z7
(18-38) AAEKTKQGVAEAAGKTKEGVL z17
(18-38) AAEKTKQGVAEAAGKTKEGVL z18
(18-38) AAEKTKQGVAEAAGKTKEGVL z19
(33-38) TKEGVL
(39-54) YVGSKTKEGVVHGVAT
(39-54) YVGSKTKEGVVHGVAT c7
(39-54) YVGSKTKEGVVHGVAT c11
(39-54) YVGSKTKEGVVHGVAT c13

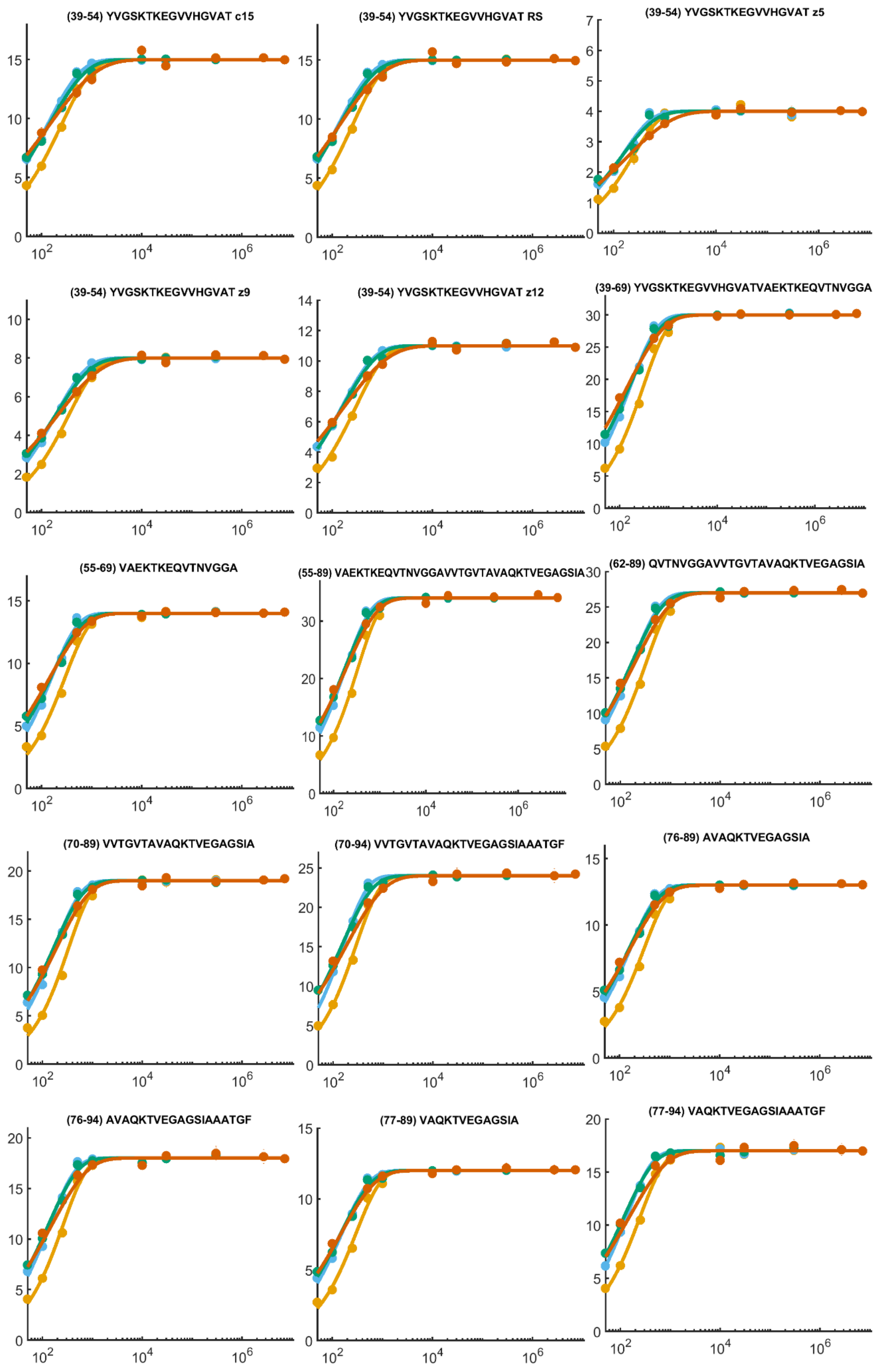
(39-54) YVGSKTKEGVVHGVAT c15
(39-54) YVGSKTKEGVVHGVAT RS
(39-54) YVGSKTKEGVVHGVAT z5
(39-54) YVGSKTKEGVVHGVAT z9
(39-54) YVGSKTKEGVVHGVAT z12
(39-69) YVGSKTKEGVVHGVATVAEKTKEQVTNVGGA
(55-69) VAEKTKEQVTNVGGA
(55-89) VAEKTKEQVTNVGGAVVTGVTAVAQKTVEGAGSIA
(62-89) QVTNVGGAVVTGVTAVAQKTVEGAGSIA
(70-89) VVTGVTAVAQKTVEGAGSIA
(70-94) VVTGVTAVAQKTVEGAGSIAAATGF
(76-89) AVAQKTVEGAGSIA
(76-94) AVAQKTVEGAGSIAAATGF
(77-89) VAQKTVEGAGSIA
(77-94) VAQKTVEGAGSIAAATGF

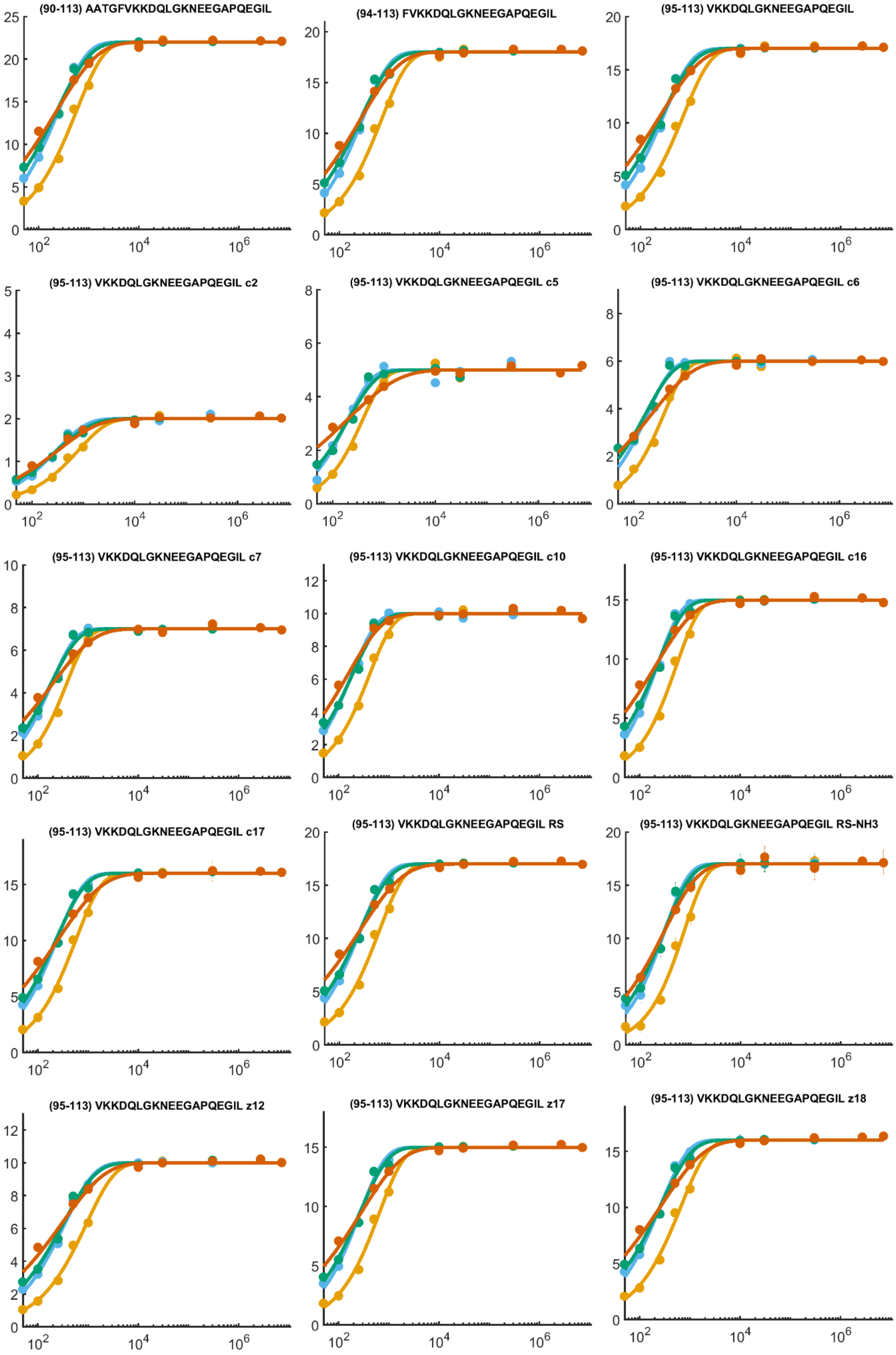
(90-113) AATGFVKKDQLGKNEEGAPQEGIL
(94-113) FVKKDQLGKNEEGAPQEGIL
(95-113) VKKDQLGKNEEGAPQEGIL
(95-113) VKKDQLGKNEEGAPQEGIL c2
(95-113) VKKDQLGKNEEGAPQEGIL c5
(95-113) VKKDQLGKNEEGAPQEGIL c6
(95-113) VKKDQLGKNEEGAPQEGIL c7
(95-113) VKKDQLGKNEEGAPQEGIL c10
(95-113) VKKDQLGKNEEGAPQEGIL c16
(95-113) VKKDQLGKNEEGAPQEGIL c17
(95-113) VKKDQLGKNEEGAPQEGIL RS
(95-113) VKKDQLGKNEEGAPQEGIL RS-NH3
(95-113) VKKDQLGKNEEGAPQEGIL z12
(95-113) VKKDQLGKNEEGAPQEGIL z17
(95-113) VKKDQLGKNEEGAPQEGIL z18

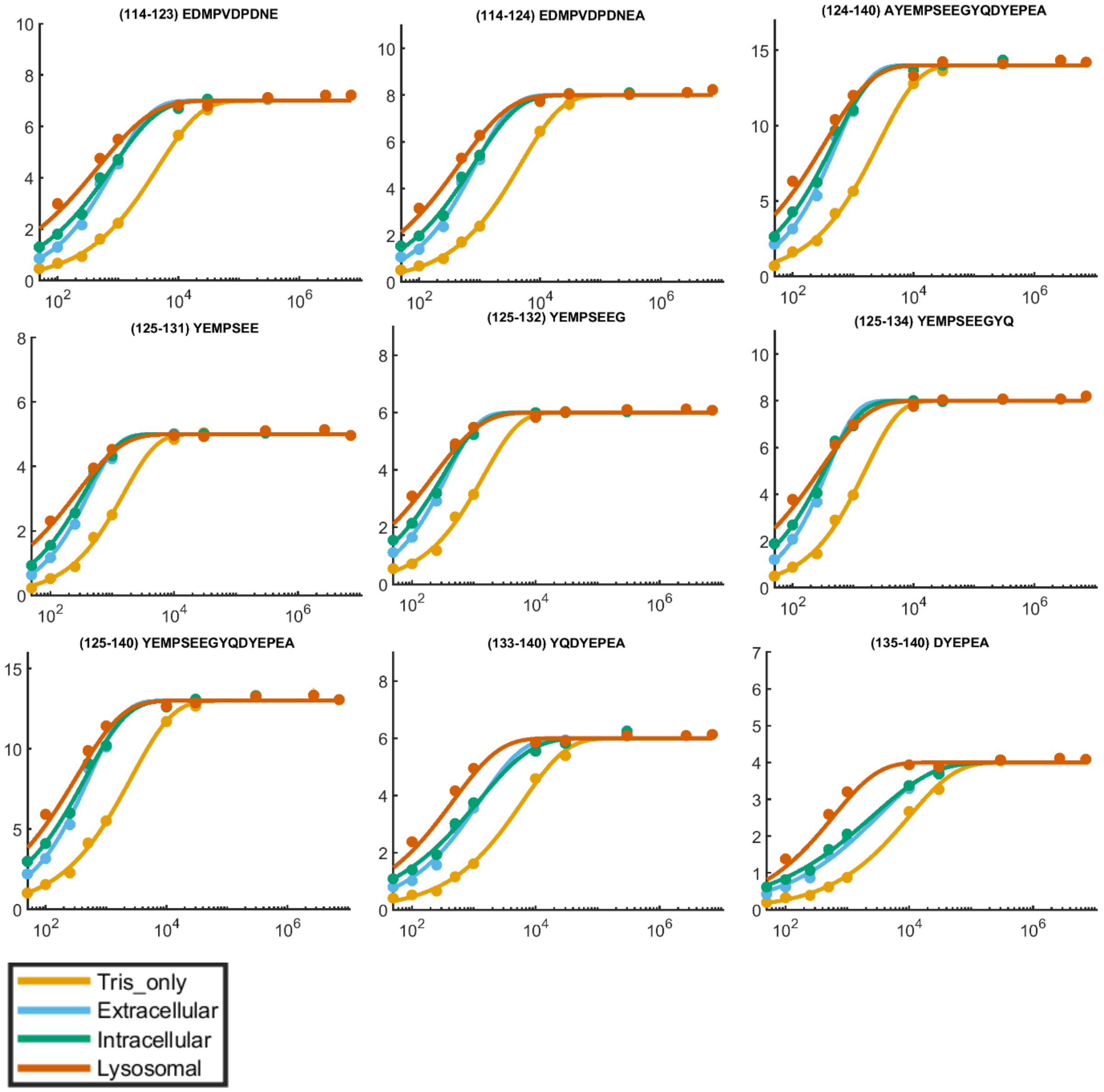

**Figure S8: Empirically adjusted deuterium uptake plots for aSyn equilibrated in Tris only, extracellular, intracellular, and lysosomal buffer conditions.** For Tris only, extracellular and intracellular states, timepoints collected were 50 ms, 100 ms, 250 ms, 500 ms, 1000 ms, 10 s, 30 s, 300 s. For lysosomal state, timepoints collected were 100 ms, 500 ms, 1000 ms, 10 s, 30 s, 300 s, 2700 s and 7200 s. Y-axis shows deuterium incorporation in Da and x-axis shows the log scale labelling times in ms (labels omitted for space). Data points are the mean of n=3 biological replicates (separate batches of aSyn protein).

**Table S1: Pearson correlation coefficients R along aSyn protein sequence (including ETD data) for correlation analyses between observed rate constant kobs from HDX-MS and nucleation lag time ($t_{lag}$) and elongation rate ($k_{agg}$) from ThT assays.**

| | Correlation coefficients | | | Correlation coefficients | | | Correlation coefficients | | | Correlation coefficients | |
|---|---|---|---|---|---|---|---|---|---|---|---|
| **Residue no.** | $k_{agg}$ | $t_{lag}$ | **Residue no.** | $k_{agg}$ | $t_{lag}$ | **Residue no.** | $k_{agg}$ | $t_{lag}$ | **Residue no.** | $k_{agg}$ | $t_{lag}$ |
| **1** | NaN | NaN | **36** | 0.669 | -0.643 | **71** | 0.710 | -0.674 | **106** | 0.748 | -0.846 |
| **2** | 0.840 | -0.667 | **37** | 0.664 | -0.636 | **72** | 0.710 | -0.674 | **107** | 0.748 | -0.846 |
| **3** | 0.840 | -0.667 | **38** | 0.664 | -0.634 | **73** | 0.710 | -0.674 | **108** | NaN | NaN |
| **4** | 0.840 | -0.667 | **39** | NaN | NaN | **74** | 0.710 | -0.674 | **109** | 0.748 | -0.846 |
| **5** | 0.846 | -0.699 | **40** | 0.627 | -0.840 | **75** | 0.710 | -0.674 | **110** | 0.748 | -0.846 |
| **6** | 0.688 | -0.408 | **41** | 0.627 | -0.840 | **76** | 0.710 | -0.674 | **111** | 0.767 | -0.832 |
| **7** | 0.688 | -0.408 | **42** | 0.627 | -0.840 | **77** | 0.763 | -0.684 | **112** | 0.780 | -0.853 |
| **8** | 0.688 | -0.408 | **43** | 0.627 | -0.840 | **78** | 0.776 | -0.687 | **113** | 0.780 | -0.853 |
| **9** | 0.747 | -0.478 | **44** | 0.627 | -0.840 | **79** | 0.776 | -0.687 | **114** | NaN | NaN |
| **10** | 0.830 | -0.728 | **45** | 0.627 | -0.840 | **80** | 0.776 | -0.687 | **115** | 0.574 | -0.938 |
| **11** | 0.830 | -0.728 | **46** | 0.716 | -0.855 | **81** | 0.776 | -0.687 | **116** | 0.574 | -0.938 |
| **12** | 0.830 | -0.728 | **47** | 0.716 | -0.855 | **82** | 0.776 | -0.687 | **117** | NaN | NaN |
| **13** | 0.830 | -0.728 | **48** | 0.716 | -0.855 | **83** | 0.776 | -0.687 | **118** | 0.574 | -0.938 |
| **14** | 0.830 | -0.728 | **49** | 0.716 | -0.855 | **84** | 0.776 | -0.687 | **119** | 0.574 | -0.938 |
| **15** | 0.830 | -0.728 | **50** | 0.720 | -0.857 | **85** | 0.776 | -0.687 | **120** | NaN | NaN |
| **16** | 0.830 | -0.728 | **51** | 0.720 | -0.857 | **86** | 0.776 | -0.687 | **121** | 0.574 | -0.938 |
| **17** | 0.830 | -0.728 | **52** | 0.747 | -0.895 | **87** | 0.776 | -0.687 | **122** | 0.574 | -0.938 |
| **18** | 0.822 | -0.805 | **53** | 0.747 | -0.895 | **88** | 0.776 | -0.687 | **123** | 0.574 | -0.938 |
| **19** | 0.591 | -0.756 | **54** | 0.732 | -0.930 | **89** | 0.776 | -0.687 | **124** | 0.582 | -0.942 |
| **20** | 0.591 | -0.756 | **55** | 0.795 | -0.859 | **90** | 0.765 | -0.548 | **125** | 0.670 | -0.945 |
| **21** | 0.591 | -0.756 | **56** | 0.800 | -0.829 | **91** | 0.776 | -0.600 | **126** | 0.672 | -0.948 |
| **22** | 0.655 | -0.726 | **57** | 0.800 | -0.829 | **92** | 0.776 | -0.600 | **127** | 0.672 | -0.948 |
| **23** | 0.655 | -0.726 | **58** | 0.800 | -0.829 | **93** | 0.776 | -0.600 | **128** | NaN | NaN |
| **24** | 0.655 | -0.726 | **59** | 0.800 | -0.829 | **94** | 0.776 | -0.600 | **129** | 0.672 | -0.948 |
| **25** | 0.767 | -0.770 | **60** | 0.800 | -0.829 | **95** | 0.773 | -0.834 | **130** | 0.672 | -0.948 |
| **26** | 0.767 | -0.770 | **61** | 0.800 | -0.829 | **96** | 0.645 | -0.769 | **131** | 0.672 | -0.948 |
| **27** | 0.767 | -0.770 | **62** | 0.800 | -0.829 | **97** | 0.634 | -0.785 | **132** | 0.678 | -0.947 |
| **28** | 0.767 | -0.770 | **63** | 0.798 | -0.805 | **98** | 0.634 | -0.785 | **133** | 0.666 | -0.937 |
| **29** | 0.767 | -0.770 | **64** | 0.798 | -0.805 | **99** | 0.634 | -0.785 | **134** | 0.602 | -0.947 |
| **30** | 0.767 | -0.770 | **65** | 0.798 | -0.805 | **100** | 0.678 | -0.778 | **135** | 0.559 | -0.928 |
| **31** | 0.771 | -0.776 | **66** | 0.798 | -0.805 | **101** | 0.700 | -0.848 | **136** | 0.454 | -0.872 |
| **32** | 0.771 | -0.776 | **67** | 0.798 | -0.805 | **102** | 0.703 | -0.895 | **137** | 0.454 | -0.872 |
| **33** | 0.766 | -0.793 | **68** | 0.798 | -0.805 | **103** | 0.703 | -0.895 | **138** | NaN | NaN |
| **34** | 0.679 | -0.653 | **69** | 0.798 | -0.805 | **104** | 0.703 | -0.895 | **139** | 0.454 | -0.872 |
| **35** | 0.679 | -0.653 | **70** | 0.731 | -0.629 | **105** | 0.748 | -0.846 | **140** | 0.454 | -0.872 |

**Table S2: Number of fibril polymorphs (n) identified per condition.**

| | Tris only | Extracellular | Intracellular | Lysosomal |
|---|---|---|---|---|
| **Polymorph p1** | 5 | 3 | 6 | 6 |
| **Polymorph p2a** | 8 | - | 11 | - |
| **Polymorph p2b** | 20 | - | 2 | - |
| **Polymorph p3a** | - | 12 | 2 | 16 |
| **Polymorph p3b** | - | - | 3 | 6 |
| **Polymorph p4** | - | - | - | - |

**Table S3: HDX-MS experimental technical details.**

| Data Set | aSyn |
|---|---|
| HDX reaction details | See Table 1 for buffer conditions. 20˚C |
| HDX time course (ms) | Tris only, Extracellular, Intracellular: 50, 100, 250, 500, 1000, 10000, 30000, 300000.<br>Lysosomal: 100, 500, 1000, 10000, 30000, 300000, 2700000, 7200000. |
| HDX control samples | Maximal deuterium uptake plateau determination |
| Back-exchange (mean / IQR) | 40.7% / 17.5% |
| # of Peptides | 30 |
| Sequence coverage | 100% |
| Peptide Redundancy | 3.79 |
| Replicates (biological or technical) | 3 (biological), 1(technical) |
| Significant differences in HDX (delta HDX > X Da) | 0.36 (95% CI, quartiles method of outlier removal) |

**Code used to calculate the Pearson correlation coefficients (MATLAB 2021b, MathWorks, USA)**

*Calculate Pearson correlation coefficients between HDX $k_{obs}$ and $t_{lag}$ and $k_{agg}$*

```
load kobs_vs_ThT_correlations_peptide_ETD.mat

%% Define colours
c = [0.4940 0.1840 0.5560; 0.6350 0.0780 0.1840; 0.9290 0.6940 0.1250;0 0.4470 
0.7410];

%% Observed rate constant vs tlag
parfor a = 1:size(all_kobs,1)

      all_R_tlag{a,1} = corrcoef(all_kobs(a,:),all_tlag(a,:),'Alpha',0.01);
      all_r_tlag(a,1) = all_R_tlag{a,1}(2);

end

%% Observed rate constant vs slope
parfor a = 1:size(all_kobs,1)

       all_R_slope{a,1} = corrcoef(all_kobs(a,:),all_slope(a,:),'Alpha',0.01);
       all_r_slope(a,1) = all_R_slope{a,1}(2);
end
```

*Draw correlation plots at each amino acid*

```
fasta = fastaread('aSN_WT.fasta');
sequence = fasta.Sequence;

%% Define colours
c = [86/255 180/255 233/255; 0/255 158/255 115/255; 213/255 94/255 0 ; 230/255 
159/255 0;
    86/255 180/255 233/255; 0/255 158/255 115/255; 213/255 94/255 0 ; 230/255 
159/255 0;
    86/255 180/255 233/255; 0/255 158/255 115/255; 213/255 94/255 0 ; 230/255 
159/255 0];

%% Observed rate constant vs tlag
parfor a = 1:size(all_kobs,1)

       if all_kobs(a,:) == 0
           continue
       end
       scatter(all_kobs(a,:),all_tlag(a,:),700,c,'filled');
       ax = gca;
       ax.YLabel.String =  'Lag time (h)'; % 'Slope (h^-1)';
       ax.XLabel.String = 'Observed rate constant (s^-1)';
       ax.LineWidth = 4;
       ax.FontSize = 30;
       %ax.XLim = [0 0.01];
       ax.YLim = [0 150];
       ax.XScale = 'linear';
       set(gcf,'Position',[493 206 1000 735]);
```

```
        % Add correlation line
        h1 = lsline(ax);
        h1.Color = 'k';
        h1.LineWidth = 5;
        h1.LineStyle = '--';

        title(sprintf('%s%d',fasta.Sequence(a),a));
        %legend([h(:)],{'Extracellular','Intracellular','Lysosomal','Tris only'});
        saveas(gcf,sprintf('kobs_vs_tlag_AA_%d.png',a));

end

%% Observed rate constant vs slope
parfor a = 1:size(all_kobs,1)

      if kobs(a,:) == 0
           continue
       end
        scatter(kobs(a,:),slope(a,:),700,c,'filled');
        ax = gca;
        ax.YLabel.String = 'Slope (h^-1)';
        ax.XLabel.String = 'Observed rate constant (s^-1)';
        ax.LineWidth = 4;
        ax.FontSize = 30;
        ax.YLim = [0.01 0.05];

        % Add correlation line
        h1 = lsline(ax);
        h1.Color = 'k';
        h1.LineWidth = 5;
        h1.LineStyle = '--';

        set(gcf,'Position',[493 206 1000 735]);
        title(sprintf('%s%d',fasta.Sequence(a),a));
        %legend([h(:)],{'Extracellular','Intracellular','Lysosomal','Tris only'});
        saveas(gcf,sprintf('kobs_vs_slope_AA_%d.png',a));

end
```
